# High regeneration-associated stress defines a distinct HCC subgroup with therapeutically exploitable vulnerabilities

**DOI:** 10.64898/2026.08.19.745832

**Authors:** Nina Desboeufs, Peter Leary, Chenhui Zhao, Sarah Kollar, Lap Kwan Chan, Lara Planas-Paz, André Fitsche, Alexander Schmidt, Fabiola Prutek, Katja R. Baumann, Simone Schneebeli, Susanne Dettwiler, Federico Donà, Reha Akpinar, Luigi M. Terracciano, Salvatore Piscuoglio, Luca Di Tommaso, Kim Wild, Laura Summermatter, Adrian Kobe, Gilbert D. Puippe, Anne-Laure Leblond, Katharina Endhardt, Charlotte K. Y. Ng, Sandro Nuciforo, Markus H. Heim, Ralph Fritsch, Chantal Pauli, Andreas E. Kremer, Massimo Lopes, Achim Weber

## Abstract

**Background:** To date, no precision oncology approach has been established for HCC. Despite the diverse underlying causes, HCC development exhibits a uniform pathophysiology characterised by chronic hyper-proliferation, resulting from hepatocyte apoptosis and compensatory liver regeneration. This chronic hyper-proliferative pressure, termed “regeneration stress”, drives genomic instability during HCC onset, yet its therapeutic potential remains poorly explored. This study aimed to identify targetable vulnerabilities tied to regeneration stress and establish clinically applicable markers for treatment stratification.

**Methods:** Weighted gene co-expression network analysis (WGCNA) was applied on external bulk RNA-seq datasets to define a LIVer REgeneration Stress Signature (LIVRESS). The signature was functionally validated using HCC patient-derived organoids (HCC-Org), and vulnerabilities were mapped using mid-throughput drug screening, single-molecule and single-cell assays, and multi-omic integration.

**Results:** High LIVRESS scores, characterised by enrichment in replication, mitotic and DNA damage repair pathways, identified a subset of HCC patients with aggressive disease and poorer survival across aetiologies. HCC-Org with high LIVRESS scores displayed exquisite sensitivity to multiple inhibitors of the checkpoint kinase ATR. Although HCC-Org models exhibited a baseline reduction in replication fork speed, sensitivity to ATR inhibitor (ATRi) was decoupled from replication fork dynamics and rather linked to intrinsic mitotic instability. ATR inhibition triggers mitotic failure and apoptosis in LIVRESS^High^ HCC-Org. This killing effect was significantly potentiated by combining ATRi with PARPi or WEE1i. Multi-omic integration identified KPNA2 as a surrogate biomarker of ATRi sensitivity.

**Conclusion:** Our findings demonstrate that a subset of HCC-Org, characterised by high liver regeneration-associated stress, is vulnerable to ATRi-based therapies. By focusing on a comprehensive regenerative stress model, we establish a framework to stratify HCC patients and implement biomarker-driven, ATR-based therapies for HCC patients with advanced disease.

**Impact and implications:** Regeneration stress is a key factor that drives genomic instability in HCC, providing a basis for the LIVRESS to identify patients dependent on ATR-mediated checkpoints. These findings reveal a conceptual shift for researchers and trialists: ATRi efficacy is decoupled from replication fork dynamics and instead leverages mitotic fragility. Practically, the LIVRESS and its IHC surrogate marker (KPNA2) offer a scalable roadmap for physicians to improve patient stratification in ATRi-based precision oncology trials. While requiring prospective validation, these results pave the way toward biomarker-driven therapies for advanced HCC.

## Introduction

HCC is a leading cause of cancer-related mortality worldwide [1]. While immune checkpoint inhibitors have transformed the treatment landscape [2], the molecular heterogeneity of HCC remains a major barrier to effective precision oncology [3–5]. While the genomic landscape of HCC is diverse [6–8], a unifying hallmark across all aetiologies is compensatory regeneration, characterised by increased hepatocyte apoptosis and concomitant proliferation [9]. This chronic proliferative pressure, termed “regeneration stress”, represents the cumulative burden of hyper-proliferation and the critical demand for genomic maintenance. While regeneration is essential for organ homeostasis, its chronic activation drives genomic instability that characterises hepatocarcinogenesis [9].

Taking advantage of the dependency on the replication and mitotic machinery to sustain hyper-proliferation, several leverage mechanisms have been explored in cancer therapy. Among them, ataxia telangiectasia and rad3-related protein (ATR) kinase inhibitors (ATRi) showed encouraging results in ongoing clinical trials [10–15]. Traditional models of ATRi efficacy focus on replication stress, broadly defined as the impairment of DNA replication [16], where ATRi triggers replication catastrophe by exhausting replication protein A (RPA) pools and inducing fork collapse [17]. However, the predictive value of genomic replication stress markers in clinical settings remains modest [18]. Furthermore, the specific contribution of the liver’s unique regenerative capacity upon DNA damage response (DDR) inhibition is poorly understood. Consequently, there is an urgent need to shift from aetiology- and genomic-based stratification toward a mechanistic framework that captures the functional vulnerabilities of the “stressed” liver.

In this study, we explored the therapeutic potential of targeting the cell cycle and DDR in mono- and combination therapy for HCC, guided by a multi-omic framework. We identified ATR as a targetable vulnerability in a subset of HCC patient-derived organoids (HCC-Org), leveraging regenerative stress, not by exacerbating replication stress, but by inducing mitotic failure and apoptosis, which was potentiated by combination with poly(ADP-ribose) polymerase inhibitor (PARPi)/WEE1i. Within the scope of this study, we demonstrated the value of a protein-based biomarker as a potential driver of clinical decision-making for future clinical trials involving ATRi monotherapy and combination therapy.

## Material and methods

A detailed description of the methodology is provided in the supplementary materials and methods.

## Results

### HCC patients with higher regeneration-associated stress exhibit poorer survival

Proliferation-associated stress, triggered by chronic liver regeneration, has been described in HCC, as evidenced by the occurrence of *γ*H2AX, a marker of DDR, and chromosomal instability in premalignant liver tissue as well as in HCC [9]. We hypothesised that the chronic burden of hyper-proliferation and genomic maintenance may manifest as a distinct gene expression network, reflecting cellular adaptations to the regeneration stress. To explore these questions, we implemented a weighted gene co-expression network analysis (WGCNA) on the differentially expressed genes (DEGs) between 15 healthy donors and 114 HCC patients (Fig. 1A). Bulk RNA-seq data of HCC tissue before treatment were obtained from a published dataset [19] and analysed as previously described by Dreyer *et al.* [20]. This unbiased approach enabled the identification of a gene module composed of functionally related and potentially co-regulated genes. Gene ontology and pathway enrichment analyses revealed that this module was enriched in genes involved in the DDR, cell cycle and division regulation and replication stress-related processes (Fig. 1B, S1A). This observation was validated in an independent HCC dataset from The Cancer Genome Atlas (TCGA – LIHC) (Fig. S1B). Among the top genes in this specific module, we identified genes involved in the ATR pathway, like *CHEK1* and *CDC25C*, acting downstream in the regulation of cell cycle checkpoints (Fig. 1C) [18]. Genes involved in both DNA damage repair and replication stress, such as *TOP2A* and *RAD51AP1,* which participate in homologous recombination and fork protection, were upregulated [21,22]. Activation of the spindle assembly checkpoint (SAC) components *BUBR1* and *CDC20*, alongside the centrosome regulator *AURKA*, key mitotic kinases, was also observed. To quantify the level of regeneration stress further, we calculated a score based on the expression of the 313 genes involved in DDR, cell cycle checkpoints and replication stress response, resulting in a regeneration stress signature for HCC (termed LIVer REgeneration Stress Signature (LIVRESS)), which we propose as a proxy for regenerative stress in the liver.

**Fig. 1:**
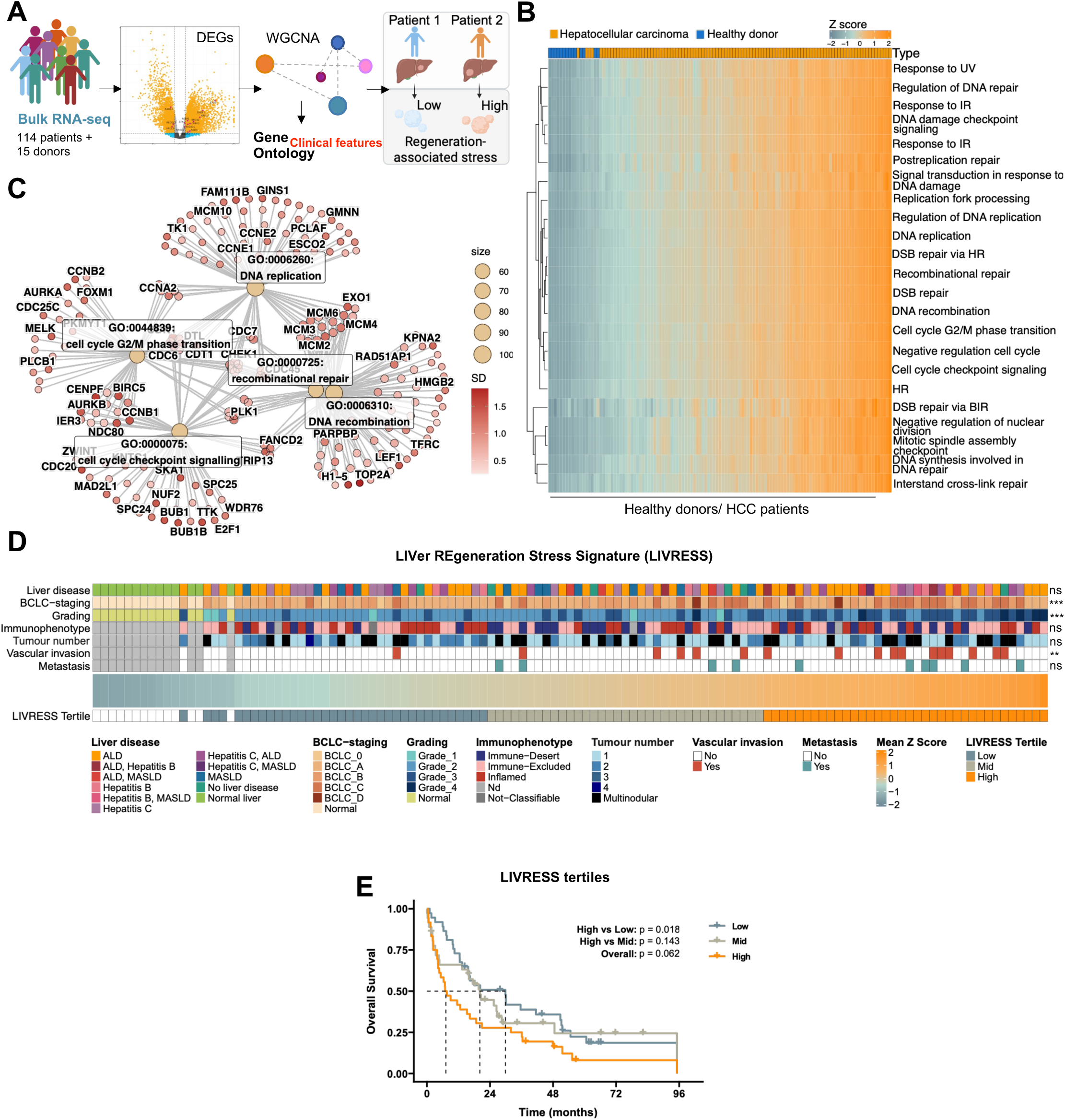
HCC patients with higher regeneration-associated stress have more aggressive disease and poorer survival. (A) Diagram depicting WGCNA implementation from healthy donors (n=15) and HCC patients (n=114), using a published dataset [19]. (B) Heatmap with clustering with LIVRESS GO terms. (C) Network plot of LIVRESS genes. (D) HCC patients’ LIVRESS score categories. Statistics: Jonckheere-Terpstra test for categorical variables, Kruskal-Wallis test for multi-level categorical variables, and Wilcoxon rank-sum test for binary variables. All *p*-values were adjusted for multiple testing using Benjamini-Hochberg test: ns > 0.05, ** ≤ 0.01, *** ≤ 0.001. (E) Kaplan Meier survival curves. Log-rank test for survival.

The ranking of healthy donor and HCC samples based on the LIVRESS score heterogeneity indicated surprisingly heterogeneous levels of regeneration stress within both HCC datasets (Fig. 1D, S1C). By analysing the clinical characteristics of the LIVRESS scores, we found no association between the signature and the underlying liver disease or the immune profile (Fig. 1D). This suggests that regeneration stress levels are independent of the disease aetiology and the immune environment. Nevertheless, we did find that the LIVRESS score was associated with the Barcelona Clinic Liver Cancer (BCLC) staging, which is a key parameter to guide treatment decisions for HCC patients [2]. Furthermore, the LIVRESS score was associated with the World Health Organisation (WHO) grading, showing that HCC patients with higher LIVRESS scores often had more poorly differentiated tumours compared to those with lower scores. Likewise, we noted an association with the LIVRESS scores and the presence of vascular invasion, indicating a more aggressive disease phenotype in HCC patients.

We then asked whether the aggressive phenotype observed in the LIVRESS^High^ HCC subset translated into differences in patient survival. To evaluate prognosis, we performed survival analysis based on LIVRESS score tertiles (low, mid, high) in the discovery cohort (Fig. 1E, S1D). While the overall log-rank test approached significance (p = 0.062), patients with high LIVRESS scores demonstrated significantly poorer survival than those in the low-score group (p = 0.018). Patient in the middle tertile did not reach statistical significance, indicating that the prognostic impact is most pronounced at the score extremes. To confirm these findings, we performed external validation in the larger, independent TCGA-LIHC dataset (Fig. S1E). In this cohort, the prognostic significance of the LIVRESS was strongly confirmed (p < 0.001).

Taken together, our findings highlighted heterogeneous regeneration stress levels in transcriptomic data from HCC patients, irrespective of the underlying liver disease or the immune environment. Moreover, high LIVRESS scores predict a more aggressive phenotype with limited treatment options and a consequent poorer prognosis, which underscores the specific medical needs of this HCC subset.

### HCC-Org recapitulate the heterogeneous regeneration-associated stress level observed in HCC patients

In light of the heterogeneity in regeneration stress levels observed in HCC, we aimed to systematically derive HCC-Org from various HCC patients to assess their regeneration stress levels, as defined by the LIVRESS, and to evaluate the response to standard-of-care and novel treatments for HCC. While several studies have successfully established HCC-Org [23–25], they all consistently relied on extracellular matrix substitutes such as Matrigel. To avoid potential interference with drug response and ensure reproducibility, we first established a new protocol to derive scaffold-free HCC-Org (Fig. 2A). In this study, we established HCC-Org from tumour resections and ultrasound-guided needle biopsies. We generated organoid models from ten HCC, obtaining a success rate of 20.8% (5/24) for tumour resections and 16.1% (5/31) for tumour biopsies, comparable to that of other studies [23,24,26]. The HCC-Org presented a typical grape-like morphology (Fig. 2B). Investigating the clinical variables of successfully established organoids, we successfully derived HCC organoids from various aetiologies, including hepatitis B and metabolic dysfunction-associated steatotic liver disease (MASLD) (Fig. 2C). These organoids were generated from moderately and poorly differentiated, but not from well-differentiated tumours. Additionally, the level of liver damage varied among patients, as evidenced by the levels of AST and ALT, as well as the presence or absence of cirrhosis. This last variance indicates that HCC-Org can be derived independently of the stage of the liver disease.

**Fig. 2.**
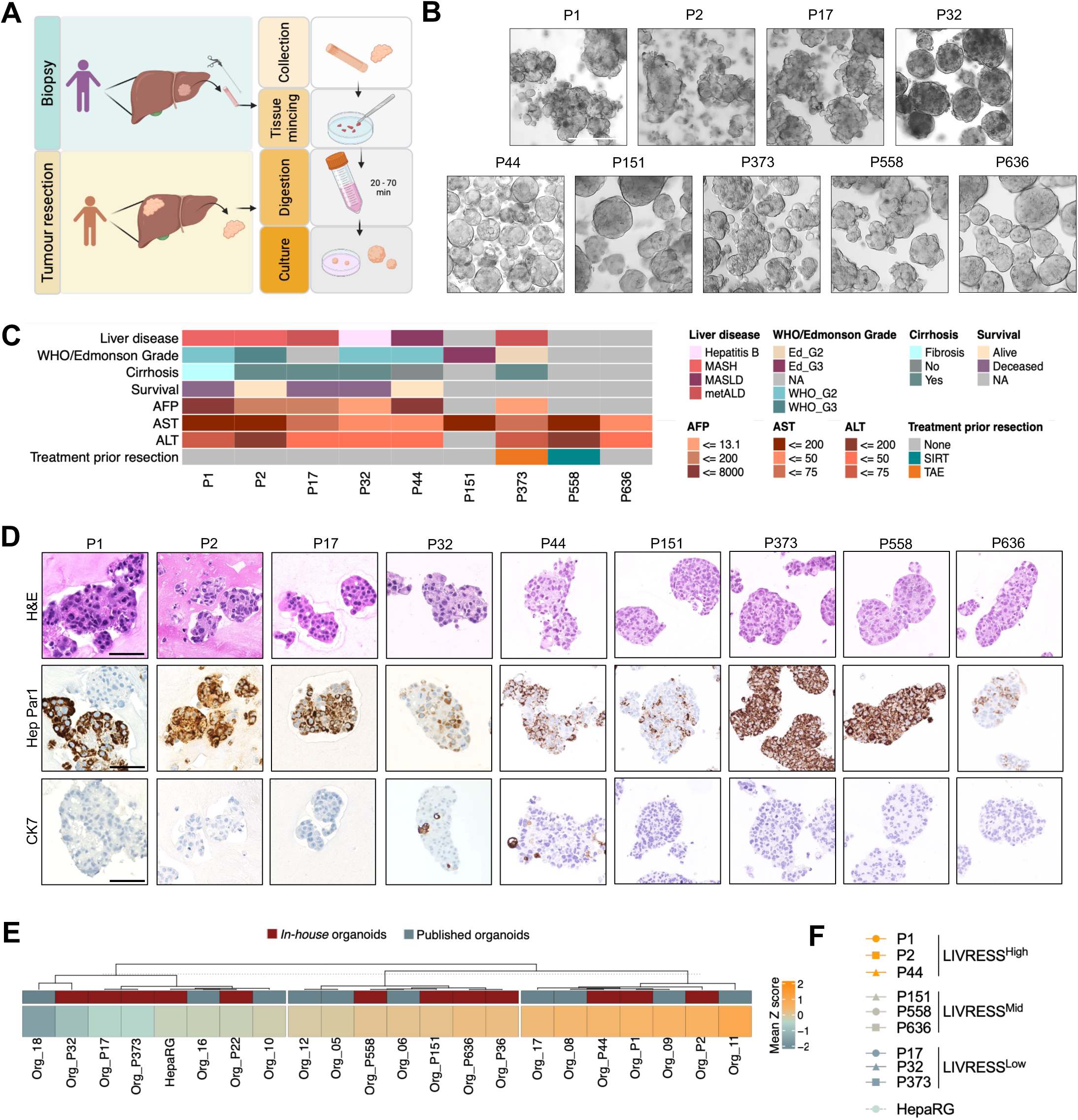
HCC-Org recapitulate the heterogeneous regeneration-associated stress landscape of clinical cohorts. (A) Schematic workflow of HCC organoids development. (B) Representative brightfield images of HCC-Org. Scale bar, 100 μm. (C) Clinical variables from patients for whom organoids were included in the study (n = 10). (D) Morphological and histological characterisation of the HCC organoids using immunohistochemical HCC and biliary markers. Scale bar, 100 μm. (E) Ranking of mean Z-score of HCC-Org from an independent cohort [24], and the *in-house* organoids based on LIVRESS score. (F) Sub-classification summary of selected HCC-Org based on LIVRESS score.

We then interrogated whether the HCC-Org, developed using our *in-house* protocol, retained the histological characteristics of the parental tumour by using diagnostic markers in histological sections (Fig. 2D, S2A). Notably, the HCC-Org preserved key morphological features of the corresponding parental tumours, as assessed by WHO grading and growth patterns (Fig. 2D, S2B). Hepatocyte paraffin 1 (Hep Par1) was consistently observed between the parental tumour and the newly derived HCC-Org, even retaining the patchy expression of Hep Par1 of the tumour from patients 1 and 17. The biliary marker cytokeratin 7 (CK7) was only found in the tumour of patients 32 and 44, where it was sparsely expressed in both the parental tumour and organoids, while it was completely absent in the other HCC-Org models. To increase the number of samples for further experiments, we also included four previously published HCC-Org (P151, P373, P558, P636), adapted them to the *in-house* medium and confirmed the HCC diagnostic (Fig. 2D) [27].

Given the heterogeneity in the LIVRESS score among HCC patients, we investigated whether HCC-Org represent different regeneration stress subsets of HCC. To this aim, we calculated the LIVRESS score in both the newly established HCC-Org and those published by Nuciforo *et al.* [24] (Fig. 2E, 2F). Notably, the ranking of the LIVRESS score of HCC-Org revealed distinct groups with low, mid and high LIVRESS scores. This observation indicates that HCC-Org retain the heterogeneous trait of the LIVRESS observed in the bulk RNA-seq data from two independent cohorts of HCC patients.

Nine HCC-Org models were selected as representative and thus appropriate for further analysis, including three with LIVRESS^Low^ scores, three with LIVRESS^Mid^ scores, and three with LIVRESS^High^ scores (Fig. 2F). Additionally, we included the HepaRG cell line in a 3D culture, a surrogate marker of non-transformed hepatocytes, which was found to be LIVRESS^Low^. The HepaRG cell line was cultured in 3D for better comparison with HCC-Org, since it was shown to enhance hepatic gene expression compared to a 2D monolayer [28].

Taken together, these results suggest that HCC-Org not only faithfully retain the histopathological features of the initial tumour but also encompass a diverse range of regeneration stress.

### ATR and WEE1 are potential therapeutic targets in HCC-Org with high regeneration-associated stress

Given that LIVRESS reflects the cumulative burden of regeneration stress, we sought to determine whether this metric could serve as a biomarker for cell cycle checkpoint, replication stress and DDR-targeting therapies. We reasoned that HCC-Org with high LIVRESS scores may be sensitive to the inhibition of pathways they rely on, imposed by their regeneration pressure. Hence, we performed a mid-throughput drug screen with a customised library that included inhibitors of the replication stress and DDR, cell cycle checkpoints, standard systemic treatments for HCC patients, and chemotherapeutics (Fig. 3A, Table S1).

**Fig. 3:**
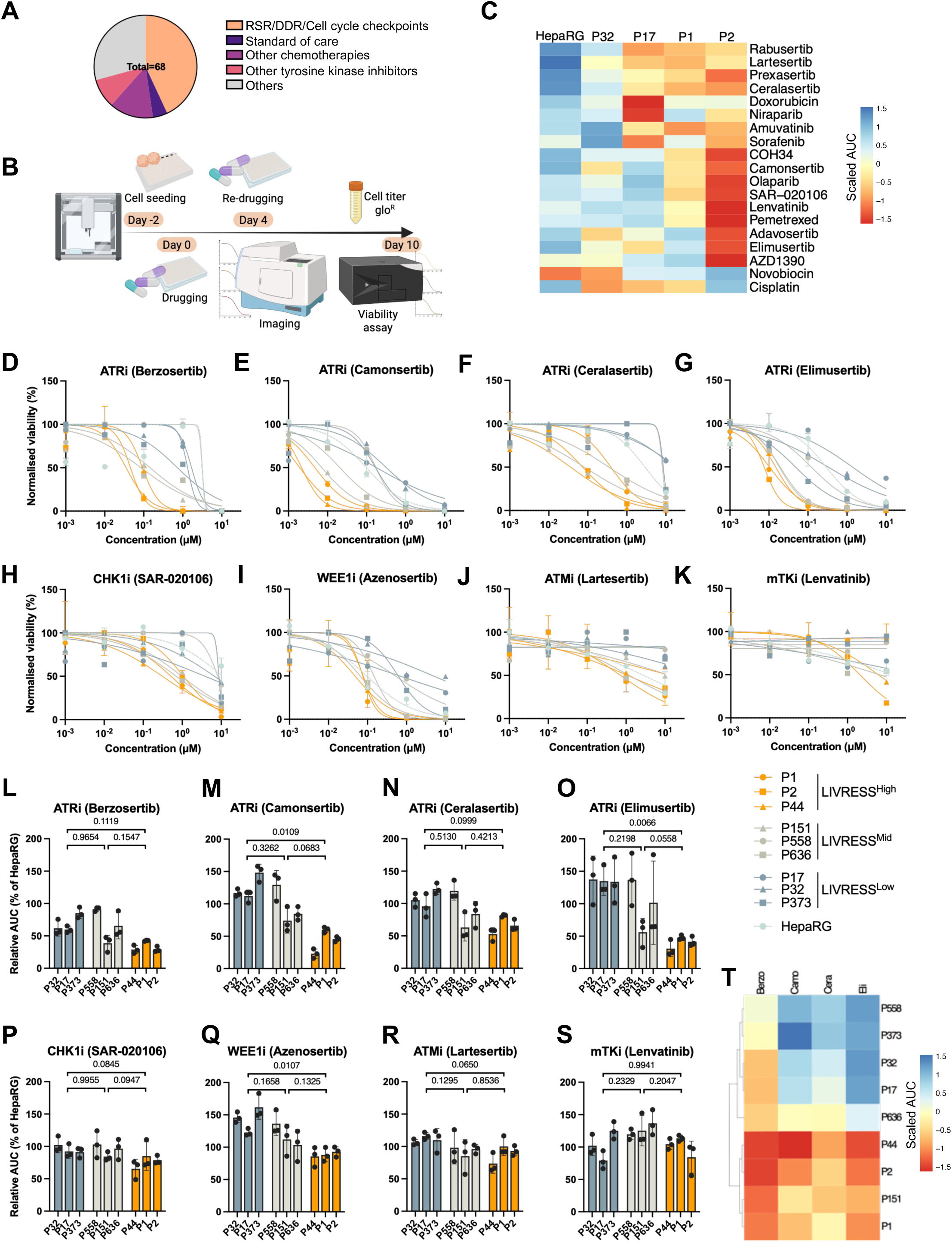
Functional screening identifies ATR as a vulnerability in high LIVRESS HCC-Org. (A) Drug screen library. (B) Mid-throughput drug screening pipeline. (C) Heatmap depicting drug response of HCC-Org and HepaRG spheroids (scaled AUC). (D-S) Viability dose-response curves (D-K) and relative AUC (L-S, normalised to HepaRG) from drug validation experiments in HCC-Org and HepaRG spheroids (normalised to DMSO control). Statistics: nested one-way ANOVA with Tukey’s multiple comparisons test. (T) Heatmap with clustering (scaled AUC) from drug validation experiments in HCC-Org. Average and SD are shown from three independent experiments. *P*-values indicated.

We performed drug screening on four HCC-Org models and HepaRG spheroids. Cells were seeded two days prior to drugging to allow for HCC-Org formation. Drug treatment was performed twice over the course of ten days, and Cell Titer-Glo^®^ was used as a surrogate readout for cell viability (Fig. 3B). Notably, we identified several compounds, including ATRi (ceralasertib, camonsertib and elimusertib), CHK1/CHK2i inhibitors (prexasertib, SAR-020106), a WEE1 inhibitor (adavosertib) and a PARG inhibitor (COH34) that preferentially affected the LIVRESS^High^ HCC-Org (Fig. 3C).

We then manually validated a selection of compounds on all ten models, following the same workflow (Fig. 3D-S, S3A-N). Notably, the three LIVRESS^High^ models exhibited higher sensitivity to ATRis compared to LIVRESS^Low^ models, which was statistically significant for camonsertib and elimusertib (Fig. 3D-G, L-O). A similar trend was observed for CHK1i within the ATR pathway, although it was not statistically significant (Fig. 3H,P, S3A,H). The WEE1i azenosertib induced a strong dose-dependent decrease in cell viability in the three LIVRESS^High^ models, while not in the LIVRESS^Low^ models (Fig. 3I,Q). Interestingly, the PARPi olaparib and niraparib did not exhibit a differential and consistent response based on the LIVRESS score (Fig. S3B,C,I,J). Furthermore, the antifolates pemetrexed and methotrexate did not demonstrate any change in cell viability in HCC-Org (Fig. S3D,E,K,L). The AUC values of the three LIVRESS^Low^ models were similar to those of the control hepatocyte model, HepaRG spheroids. In comparison, the three LIVRESS^Mid^ models showed an attenuated response compared to the high-score models for most of the targeted agents (Fig. 3D-S, S3A-N).

Since ATR is a member of the phosphatidylinositol 3-kinase-related kinases (PIKKs) family, such as ataxia-telangiectasia mutated protein (ATM), we also examined the response to ATM inhibitors. In contrast to ATRi, we did not observe any differential response to ATMi within the LIVRESS score for both ATMi (lartesertib and AZD1390), highlighting the specific response of ATR inhibition (Fig. 3J,R, S3F,M). Similarly, there was no differential response observed for the two multi-tyrosine kinase inhibitors (mTKi, sorafenib and lenvatinib), which are part of the systemic treatment for HCC patients (Fig. 3K,S, S3G,N). To assess the association of the LIVRESS with ATRi response, we performed hierarchical clustering on the AUC from the validation experiments across the nine HCC-Org models (Fig. 3T). Both extreme tertiles (LIVRESS^Low^ and LIVRESS^High^) samples clustered separately, while the LIVRESS^Mid^ models were distributed within both clusters.

Taken together, these results suggest that ATRi and WEE1i are promising targets for systemic therapy in a subset of HCC-Org, indicated by their levels of regeneration stress.

### ATRi sensitivity is decoupled from replication fork dynamics and linked to mitotic failure

We then investigated whether the association observed between the regeneration stress score (LIVRESS) and the sensitivity to the ATRi reflects specific features of these tumours in terms of DNA replication dynamics, such as origin firing, fork and mitotic progression, either before or in response to these treatments. We first compared replication fork progression at the single-nucleotide level in HCC-Org and HepaRG spheroids using halogenated nucleotide labelling, along with a DNA fibre spreading assay optimised for 3D models (Fig. 4A, see supplementary materials and methods). Eight out of nine HCC-Org models demonstrated a reduction in fork speed, compared to HepaRG spheroids, albeit only statistically significant for HCC-Org P2 (Fig. S4A). Also, no significant difference in fork speed was observed within LIVRESS groups (Fig. 4B,C).

**Fig. 4.**
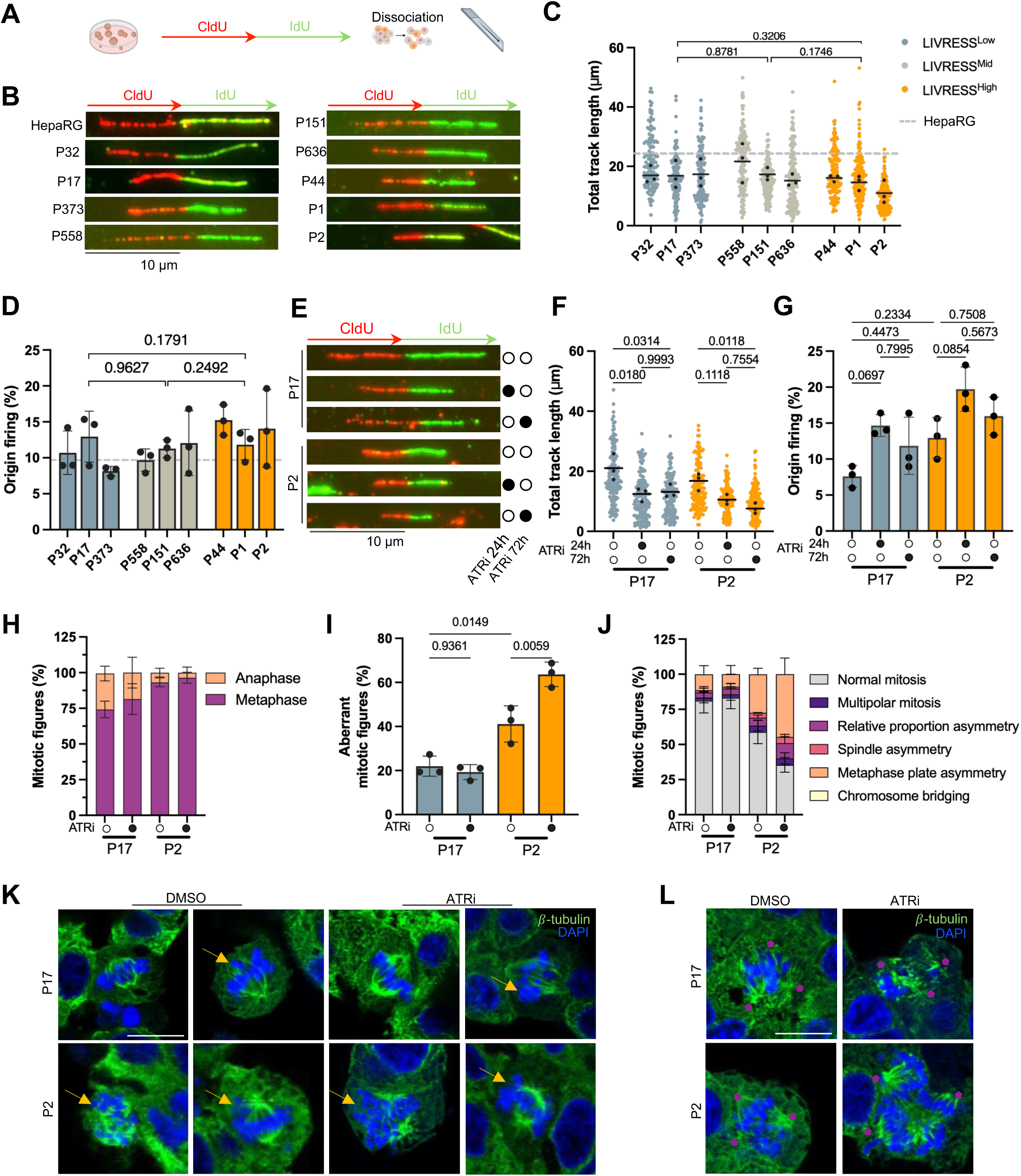
ATRi sensitivity is decoupled from replication fork dynamics and is linked to mitotic fragility. (A) DNA fibre assay schematic. (B–G) Representative images, total length and origin firing of DNA fibres in untreated HCC-Org/HepaRG (B-D) and HCC-Org ATRi (ceralasertib, 500 nM) (E-G). Scale bar, 10 μm. (H–L) Mitotic analysis via immunofluorescence (72h ± ATRi). Graphs show anaphase-to-metaphase ratios (H) and aberrant mitotic figures (I, J). Representative images of asymmetric (K) and multipolar (L) metaphase plates; arrows: unaligned chromosomes; asterisks: spindle poles. Scale bar, 10 μm. Statistics: Nested (C,D) or one-way ANOVA with Tukey’s test (F,G,I). *P*-values indicated.

We then asked whether the reduced fork speed of HCC-Org might be linked with changes in origin firing, as a reduced fork speed could indicate deregulated origin firing or be compensated by firing of nearby dormant origins to avoid under-replication (Fig. 4D) [29]. However, no difference was observed within LIVRESS tertiles in the frequency of origin firing. Overall, these data suggest that HCC-Org may experience slower endogenous fork progression, although this is not associated with specific defects in origin firing.

We then investigated the early response of HCC-Org to ATRi in terms of fork progression and origin firing. We first assessed the fork speed upon ATRi treatment in one LIVRESS^Low^ and one LIVRESS^High^ model (Fig. 4E,F). We observed a reduction in fork speed in both models following ATRi treatment. As inhibiting ATR typically leads to unscheduled origin firing by unrestraining ATR-mediated control of origin firing [30,31], we then tested the impact of ATRi on origin firing in our experimental system (Fig. 4G). As expected, ATRi induced increased origin firing in both LIVRESS^Low^ and LIVRESS^High^ HCC-Org models. Consistent with the lack of response to ATMi treatment (Fig. 3J,R, S3F,M), we also did not observe any detectable effects of ATMi on fork progression in HCC-Org (Fig. S4B,C). Overall, the observed differential response to ATRi did not reflect specific defects in replication dynamics.

Among the extensive palette of ATR functions, we then investigated the potential impact of ATRi on mitotic progression by mitotic figures analysis using *β*-tubulin immunostaining. Intriguingly, both HCC-Org models showed few anaphases compared to metaphases, suggesting a metaphase-to-anaphase delay (Fig. 4H). Additionally, HCC-Org P2 seems to present a higher mitotic arrest baseline, consistent with the enrichment of mitotic checkpoint signalling observed at the transcriptomic level (Fig. 1C). Furthermore, we observed an intrinsic higher level of aberrant mitotic figures in LIVRESS^High^ HCC-Org P2 compared to LIVRESS^Low^ HCC-Org P17 (Fig. 4I-L). The presence of aberrant mitotic figures was further and selectively exacerbated in HCC-Org P2 by ATRi. Among mitotic aberrancies, we primarily observed asymmetric metaphase plate, followed by asymmetric metaphase plate proportion, asymmetric spindle and multipolar mitosis (Fig. 4J-L).

Collectively, these results suggest that sensitivity to ATRi is decoupled from canonical replication fork dynamics. Instead, ATRi leverages cell division stress in the LIVRESS^High^ HCC-Org to trigger mitotic failure.

### ATRi demonstrates synergistic effects combined with PARPi and WEE1i in HCC organoids with high regeneration-associated stress

Several clinical trials investigating combination therapy with ATR and PARPi have demonstrated efficacy in various cancers, including *BRCA1/2*-mutated and even DNA-repair proficient cancers [32–34]. However, to our knowledge, none of these trials has included patients with HCC.

Therefore, building on the promising data we obtained with ATRi, we explored the potential of ATRi-based combination therapies in LIVRESS^High^ HCC-Org, explicitly focusing on the ATRi ceralasertib. The drug combination pipeline consisted of seeding the cells, followed by drug administration, either monotherapy or combination therapy, two days later. The cell viability was measured ten days after drugging, defining synergies when the four synergy scoring models, *i.e.*, BLISS, ZIP, LOEWE and HSA, yielded scores above ten at specific drug combination doses.

We first combined the ATRi ceralasertib with the PARPi olaparib in three LIVRESS^High^ HCC-Org to potentially increase the load of genomic instability (Fig. 5A,B) [35]. ATRi combination with PARPi showed a synergistic effect in the two HCC-Org P1 and P2, even at ATRi concentrations below 100 nM, whereas only additive effects were observed in HCC-Org P44, restricted to higher doses. Since WEE1 acts as a safeguard against mitotic catastrophe, we then evaluated the combination of the ATRi ceralasertib with WEE1i azenosertib to exacerbate mitotic failure further (Fig. 5C,D) [36,37]. Notably, this combination showed potent and uniform synergism across all three LIVRESS^High^ HCC-Org. Even nanomolar concentrations outperformed both monotherapies, highlighting it as the most promising combination therapy tested so far in the study.

**Fig. 5.**
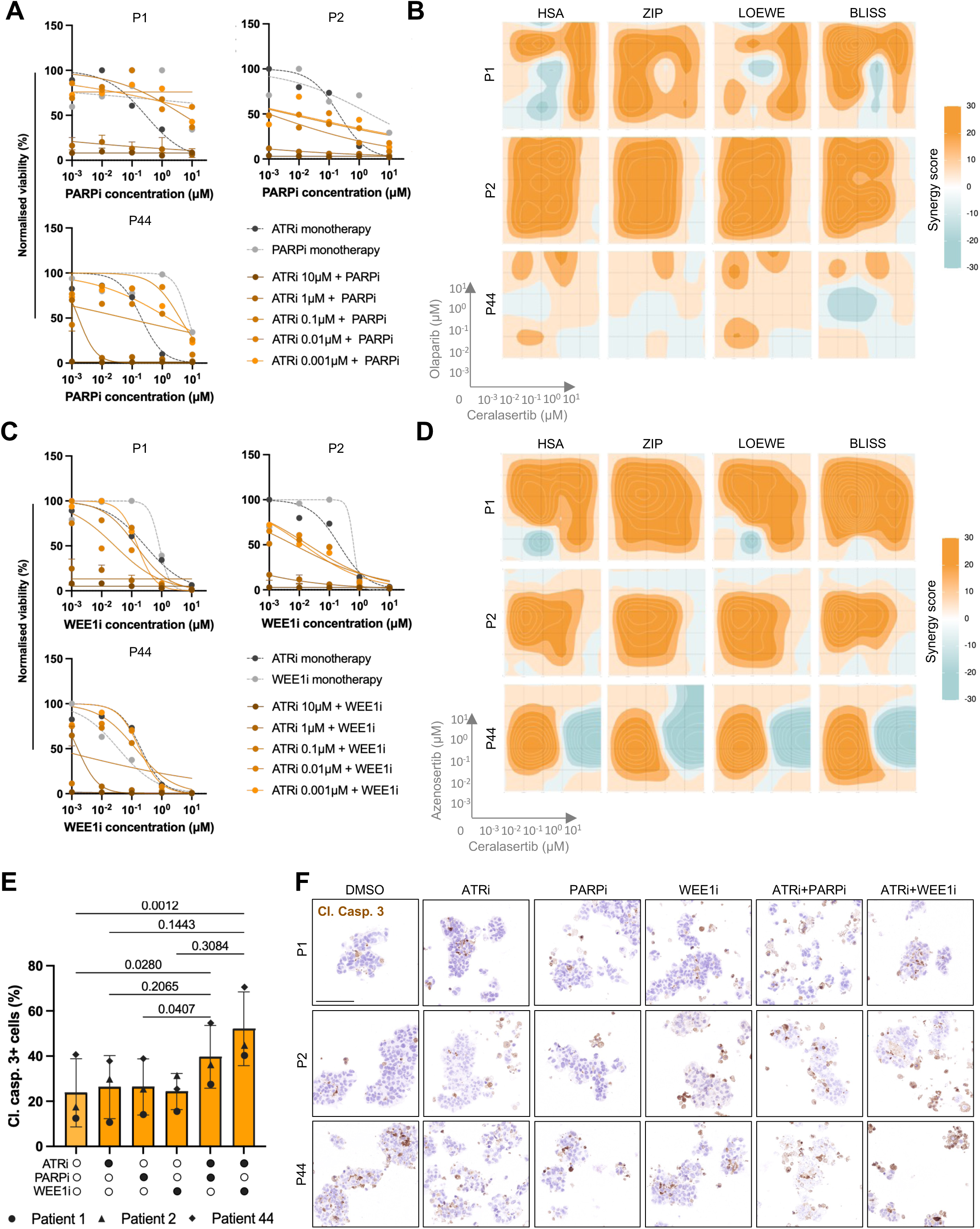
ATRi synergises with PARPi/WEE1i in LIVRESS-high HCC-Org. (A, C) Viability dose-response curves for ATRi (ceralasertib) combined with PARPi (olaparib) or WEE1i (azenosertib) in LIVRESS-high HCC-Org (10-day treatment). (B, D) Synergy landscapes across four models (HSA, ZIP, LOEWE, BLISS) visualised via 2D contour plots. (E, F) Cleaved caspase 3 (Cl. Casp. 3) IHC quantification and representative images ± monotherapy or combination (72h). ATRi (250 nM), PARPi (1 *μ*M), WEE1i (200 nM). Scale bar, 100 μm. Data represent means (n >700 cells/condition). Statistics: One-way RM ANOVA with Tukey’s test. *P*-values indicated.

To determine whether the synergistic reduction in cell viability observed in the CTG assays was driven by induced cell death, we next evaluated the pro-apoptotic effects of both combination therapies, using cleaved caspase 3 immunostaining. Already after 72h of treatment, ATRi combined with PARPi/WEE1i showed a significant increase in apoptotic cells compared to the control-treated condition in the three LIVRESS^High^ HCC-Org, while this effect was not observable in the monotherapy treatment (Fig. 5E,F).

Altogether, we observed strong efficacy of the ATRi in combination therapy, either with PARPi or WEE1i in LIVRESS^High^ HCC-Org, eliciting apoptosis.

### Integrating proteomic and phosphoproteomic identifies prototype markers for patient stratification

Building upon the transcriptomic evidence of ATRi sensitivity, we performed integrated proteomic and phosphoproteomic profiling of the LIVRESS to bridge the gap between high-throughput sequencing and clinically applicable diagnostics. This transition sought to identify protein-based biomarkers that can be easily implemented into everyday practice using IHC, thereby circumventing the inherent challenges associated with mRNA-based signatures in routine clinical use.

We first investigated if the transcriptome signature (LIVRESS) was associated with proteomic and phosphoproteomic traits by integrating the published HCC patient dataset from Ng and colleagues [19]. Principal component analysis demonstrated that HCC segregated according to the LIVRESS score, although with high variability, particularly pronounced in the LIVRESS^High^ group at the transcriptome, proteome and phosphoproteome levels (Fig. 6A, S5A). To determine the extent to which the transcriptomic signature matches functional protein activity, we correlated the LIVRESS mRNA signature and its corresponding proteomic (109/313) and phosphoproteomic (77/313) counterparts (Fig. S5B). A weak correlation was observed across both modalities (0.277 and −0.240, respectively), suggesting significant post-transcriptional and post-translational divergence. We then identified dysregulated proteins and phosphorylation sites in LIVRESS^High^ tumours by performing differential expression analyses based on the LIVRESS score. 49 and 188 proteins were up- and down-regulated in LIVRESS^High^ tumours, respectively, and 167 and 282 phosphosites were hyper- and hypophosphorylated, respectively (Fig. 6B). Notably, pathway enrichment analysis revealed that most of the upregulated proteins and hyperphosphorylated proteins in LIVRESS^High^ tumours were involved in cell cycle checkpoints, mitotic progression, DNA repair and ATR activation pathways (Fig. 6C). In contrast, downregulated proteins and hypophosphorylated proteins in LIVRESS^High^ tumours were involved in metabolism of amino acids, steroids, bile acids, xenobiotic and fatty acids pathways.

**Fig. 6.**
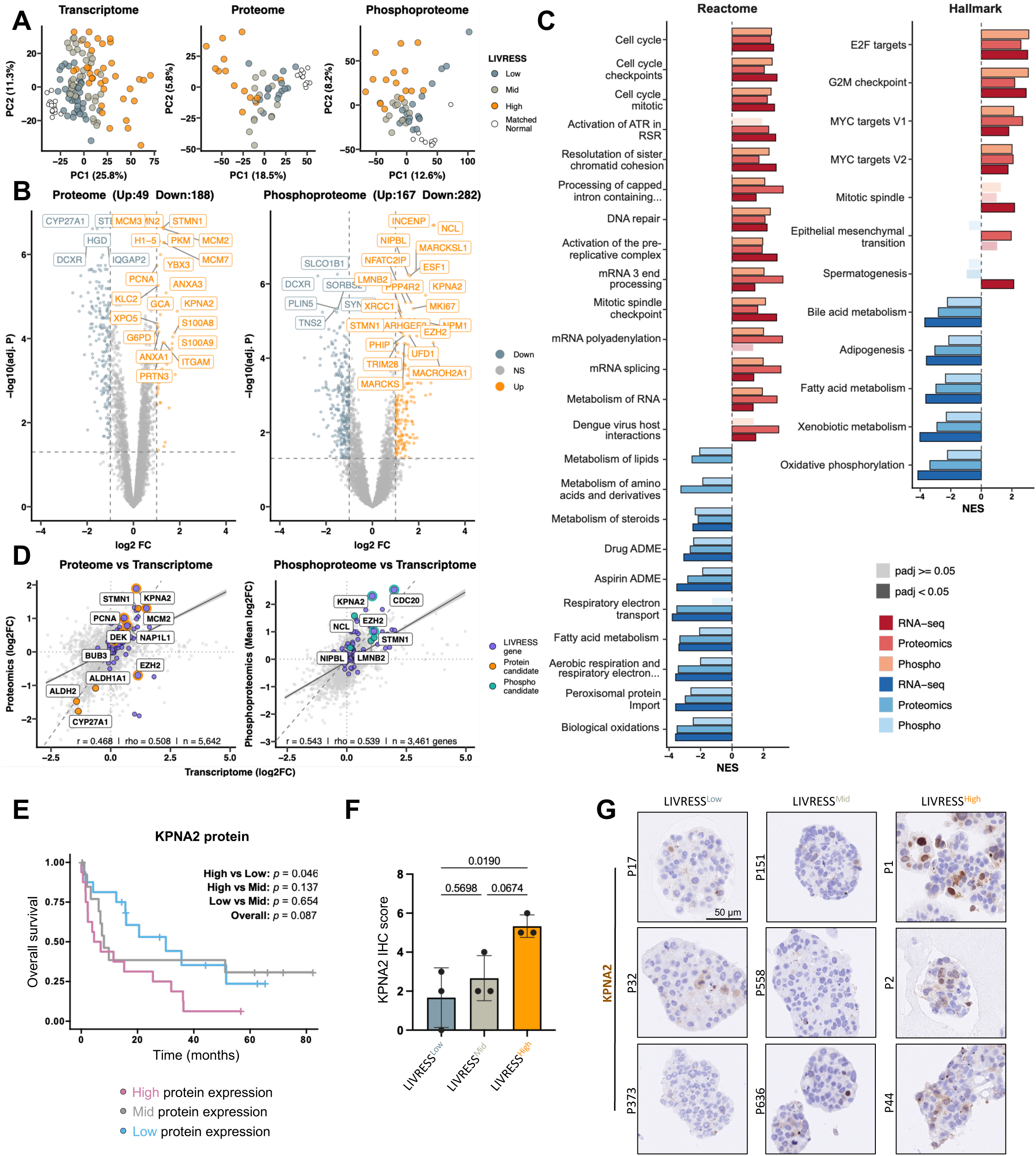
Multi-omic profiling identifies LIVRESS surrogate markers linked to poor HCC prognosis. (A) PCA of transcriptome (n=124), proteome (n=54), and phosphoproteome (n=55) in HCC and normal liver biopsies stratified by LIVRESS. (B, C) Volcano plots of LIVRESS-correlated proteins/phosphosites and top-enriched pathways in LIVRESS-high tumours. (D) Cross-omic correlations between transcriptomic, proteomic (n = 5644), and phosphoproteomic (n = 3462) profiles. (E) Kaplan-Meier survival curves. (F, G) IHC semi-quantification (F) and representative images (G) of KPNA2 on HCC-Org. Statistics: Pearson (r) and Spearman (ρ) correlations; Log-rank test for survival. One-way ANOVA with Tukey’s test (F). *P*-values indicated, dataset [19].

We then investigated proteins and phosphorylation sites associated with the LIVRESS mRNA signature as potential surrogate biomarkers of ATRi sensitivity. During selection, we prioritised IHC-compatible candidates to facilitate seamless integration into routine clinical workflows. Among the top correlated proteins, we found an enrichment of STMN1, KPNA2, and DEK, all involved in cell cycle regulation (Fig. 6D) [38]. Metabolic enzymes, including the aldehyde dehydrogenase family (ALDH1A1, ALDH2) and cytochrome P 450 enzyme (CYP27A1), were also correlated with the LIVRESS. In the phosphoproteomic landscape, STMN1 (s38) and EZH2 (thr345) emerged as top candidates with available antibodies for IHC. Of note, several of these candidates were core components of the LIVRESS.

We then focus on the top proteomic candidate, KPNA2. Kaplan-Meier analysis demonstrated that high expression of KPNA2 was associated with overall survival: patients with high expression demonstrated a worse prognosis than patients with low expression (Fig. 6E, S5C). Patients with intermediate expression profiles exhibited a mixed clinical phenotype. This was consistent with the previously observed prognosis of the LIVRESS, confirming disease aggressiveness in this HCC subset. To validate the potential of KPNA2 as a surrogate biomarker of the LIVRESS, we performed IHC for KPNA2 in an HCC tissue microarray (TMA) and HCC-Org models (Fig. 6F,G, S5D). Among patient specimens, KPNA2 demonstrated marked inter-patient heterogeneity by semi-quantification of KPNA2 (KPNA2 IHC score): nuclear expression ranged from occasional weak to strong nuclear positivity to a complete absence of staining. Crucially, KPNA2 nuclear localisation mirrored the LIVRESS stratification in HCC-Org with higher KPNA2 IHC scores in LIVRESS^High^ compared to LIVRESS^Low^ models (Fig. 6F). Models with intermediate LIVRESS scores exhibited mixed results, consistent with the transcriptomic and functional observations (Fig. 6G). LIVRESS^High^ models displayed a characteristic patchy, moderate to strong nuclear staining, whereas expression was absent from the nucleus in LIVRESS^Low^ models.

Collectively, these data identify KPNA2 immunohistochemical reactivity as a surrogate for regeneration-associated stress and a promising biomarker for identifying patients likely to benefit from ATRi therapy.

## Discussion

Precision oncology for HCC remains hindered by high inter- and intra-patient heterogeneity [4,7]. While chronic hyper-proliferation is a hallmark of hepatocarcinogenesis [9], its therapeutic potential remains under-explored. In this study, we shift our focus from observing this hallmark to exploiting the regenerative stress due to chronic hyper-proliferation for therapeutic purposes. By establishing the LIVer REgeneration Stress Signature (LIVRESS), we identified a high-risk patient subgroup, independent of the aetiology, characterised by a strong dependency on mitotic checkpoints to manage intrinsic genomic instability.

Expanding on the paradigm we have previously established [9], our findings translate the observable phenomenon of regeneration-induced damage into a quantifiable transcriptomic landscape. While previous studies defined replication stress signatures through different methodologies [20,28,39,40], LIVRESS is tailored to the liver’s unique regenerative capacity, not only focusing on replication stress but also broadening to DDR and mitotic stress. Elevated LIVRESS scores identify aggressive disease, regardless of the aetiology, highlighting a clinical subgroup to prioritise for novel therapeutic strategies.

Moreover, by mapping this broader stress landscape, we identify the specific vulnerabilities of this subgroup in HCC-Org and provide a surrogate biomarker to facilitate its implementation in clinical settings. Crucially, matched normal tissue in the TCGA-LIHC validation dataset exhibited low LIVRESS scores, suggesting that regeneration-associated stress in HCC may evolve to become quantitatively distinct from that found in chronic liver disease (CLD) [9,41]. This divergence provides a potential therapeutic window, which is vital in HCC management due to the risk of inducing hepatic decompensation through off-target effects on the underlying diseased liver [42].

Our findings provide a mechanistic refinement to current models of ATRi efficacy. While traditional models focus on replication catastrophe and fork collapse [17,43], the modest predictive power of genomic replication stress markers in clinical trials suggests a more nuanced landscape [18,44]. A central finding of our study is the decoupling of ATRi sensitivity from replication fork dynamics in HCC. Unlike previous approaches, which relied on proxies of DNA damage [39], DNA fibre spreading assays demonstrate that ATRi response is independent of changes in fork progression or origin firing. Instead, the baseline mitotic instability and high efficacy of WEEi combinations in LIVRESS^High^ models suggest that ATRi sensitivity is linked to a dependency on the G2/M checkpoint [40,45]. We postulate that the liver’s unique regenerative capacity fundamentally alters the cellular response to checkpoint inhibition; by bypassing ATR-mediated safety brakes to mitotic entry, high-LIVRESS cells are forced into mitotic failure. While combining WEE1i with conventional chemotherapy has been reported [40], the integration of DDR, replication, and mitotic stress provided by the LIVRESS identified targeted combinations of ATRi and WEE1i/PARPi. These combinations significantly enhance killing capacity in LIVRESS^High^ models, most likely by increasing the load of genomic instability or further deregulating cell cycle control, respectively [35,37].

To facilitate clinical implementation, we identified a protein-based surrogate biomarker, KPNA2, suitable for IHC-based diagnostic workflows. KPNA2 is a central karyopherin responsible for the nuclear import of the key proteins involved in DNA repair and cell cycle progression in HCC [46]. Furthermore, nuclear KPNA2 expression has been described as a prognostic marker in HCC, linked to advanced disease and tumour progression [47,48]. Here, we demonstrate that nuclear KPNA2 accumulation serves as a potential functional readout, which may be required to sustain the high regeneration stress load. By bridging the gap between transcriptomic profiling and routine pathology, KPNA2 IHC offers a scalable roadmap to identify patients who may benefit from ATRi-based therapy.

While *ex vivo* models provide a robust mechanistic foundation, they do not fully recapitulate the physiological impact of the cirrhotic stroma or the immune microenvironment. This impact may alter the tumour response to ATRi-based treatment but also challenge the management of haematological and gastrointestinal toxicities in the lesioned liver [49]. Consequently, while the LIVRESS with its surrogate biomarker demonstrated strong predictive value across a small heterogeneous HCC-Org cohort, prospective clinical validation of our findings will be needed to confirm their utility in guiding therapeutic decisions in clinical settings. Nevertheless, by establishing this framework, we offer a mechanistic-based strategy for patient stratification that goes beyond traditional aetiological boundaries.

In conclusion, our work refines regeneration-associated stress as a key driver of genomic instability and a primary therapeutic vulnerability in HCC. By shifting focus from the initial source of liver injury to the transcriptional and proteomic landscape of the adaptive response, we uncover a unique Achilles’ heel in high-stress HCC-Org: a dependency on ATR to manage regeneration-associated stress. This study not only refines the biological framework of ATRi efficacy in HCC but also offers a clinical roadmap for future biomarker-driven, precision oncology trials in advanced HCC.

## Abbreviations

ATM: ataxia-telangiectasia mutated protein
ATR: ataxia telangiectasia and rad3 related protein
ATRi: ATR inhibitor
BCLC: Barcelona Clinic Liver Cancer
BIR: break-induced replication
CK7: cytokeratin 7
Cl. Casp. 3: cleaved-caspase 3
CLD: chronic liver disease
CldU: 5-chloro-2’-deoxyuridine
DDR: DNA damage response
DEGs: differentially expressed genes
DSBs: DNA double-strand breaks
GO: gene ontology
HCC-Org: HCC patient-derived organoid
Hep Par1: hepatocyte paraffin 1
HR: homologous recombination
ICL: interstrand cross-link
IdU: 5’-iodo-2’-deoxyuridine
MASLD: metabolic dysfunction-associated steatotic liver disease
mTKi: multi-tyrosine kinase inhibitors
PARPi: poly(ADP-ribose) polymerase inhibitor
PIKKs: phosphatidylinositol 3-kinase-related kinases
RPA: replication protein A
SAC: spindle assembly checkpoint
TCGA: the cancer genome atlas
TMA: tissue microarray
TPM: transcripts per million
UV: ultraviolet
WGCNA: weighted gene co-expression network analysis
WHO: World Health Organisation.

## Acknowledgments

We thank all the patients who participated in this study, our colleagues Dr Jana Krietsch, Dr Marc Healy and Dr Gonzalez Acosta for their valuable input throughout the project, as well as the Gastroenterology and Hepatology, Interventional Radiology teams of University Hospital Zurich for their support in the development of HCC-Org. Imaging was performed with support from the Centre for Microscopy and Image Analysis, University of Zurich. Sequencing was performed at the Functional Genomics Centre Zurich (FGCZ) of the University of Zurich and ETH Zurich. We want to acknowledge the use of Grammarly and Gemini (Google) to edit the grammar and enhance the clarity of sentence formulation.

## Authors’ contributions

Contribution to conception and design: N.D., A.W., M.L.; Supervision: M.L., A.W.; Acquisition of data and/or analysis and/or interpretation of data: N.D., P.L., C.Z., S.K., L.P., K.W., A.W., L.D.T.; Patient recruitment and/or biopsy collection: A.E.K., N.D., A.K., G.D.P., A.L., K.E.; HCC-Org development and/or culture: N.D., C.Z., C.P., L.S., F.D., R.A.; Drug screening design: C.P., L.P.; Immunohistochemistry staining: A.F., A.S., S.S., F.P., S.D.; Conceptual input: C.P., L.P., A.E.K., R.F., L.M.T., L.D.T., S.P., C.K.Y.N., S.N., M.H.H.; Draft of the manuscript: N.D.; Revising the manuscript: N.D., A.W., M.L.

## References

[1] Bray F, Laversanne M, Sung H, et al. Global cancer statistics 2022: GLOBOCAN estimates of incidence and mortality worldwide for 36 cancers in 185 countries. CA Cancer J Clin 2024;74:229–63. 10.3322/caac.21834.

[2] Reig M, Forner A, Rimola J, et al. BCLC strategy for prognosis prediction and treatment recommendation: The 2022 update. J Hepatol 2022;76:681–93. 10.1016/j.jhep.2021.11.018.

[3] Sangro B, Argemi J, Ronot M, et al. EASL Clinical Practice Guidelines on the management of hepatocellular carcinoma. J Hepatol 2024.

[4] Yang X, Yang C, Zhang S, et al. Precision treatment in advanced hepatocellular carcinoma. Cancer Cell 2024;42:180–97. 10.1016/j.ccell.2024.01.007.

[5] Mauro E, de Castro T, Zeitlhoefler M, et al. Hepatocellular carcinoma: Epidemiology, diagnosis and treatment. JHEP Reports 2025;7. 10.1016/j.jhepr.2025.101571.

[6] Zucman-Rossi J, Villanueva A, Nault JC, et al. Genetic Landscape and Biomarkers of Hepatocellular Carcinoma. Gastroenterology 2015;149:1226–1239.e4. 10.1053/j.gastro.2015.05.061.

[7] Friemel J, Rechsteiner M, Frick L, et al. Intratumor heterogeneity in hepatocellular carcinoma. Clinical Cancer Research 2015;21:1951–61.

[8] Wu Y, Liu Z, Xu X. Molecular subtyping of hepatocellular carcinoma: A step toward precision medicine. Cancer Commun 2020;40:681–93.

[9] Boege Y, Malehmir M, Healy ME, et al. A Dual Role of Caspase-8 in Triggering and Sensing Proliferation-Associated DNA Damage, a Key Determinant of Liver Cancer Development. Cancer Cell 2017;32:342–359.e10. 10.1016/j.ccell.2017.08.010.

[10] da Costa AABA, Chowdhury D, Shapiro GI, et al. Targeting replication stress in cancer therapy. Nat Rev Drug Discov 2023;22:38–58.

[11] Cybulla E, Vindigni A. Leveraging the replication stress response to optimise cancer therapy. Nat Rev Cancer 2023;23:6–24.

[12] Cote GM, Kochupurakkal BS, Do K, et al. A Translational Study of the ATR Inhibitor Berzosertib as Monotherapy in Four Molecularly Defined Cohorts of Advanced Solid Tumours. Clinical Cancer Research 2025;31:35–44.

[13] Dillon MT, Guevara J, Mohammed K, et al. Durable responses to ATR inhibition with ceralasertib in tumours with genomic defects and high inflammation. J Clin Invest 2024;134.

[14] Carneiro BA, Rosen E, Fontana E, et al. 619MO Camonsertib (cam) monotherapy in patients (pts) with advanced cancers harbouring ATM loss-of-function (LoF). Annals of Oncology 2024;35:S496.

[15] Yap TA, Fontana E, Lee EK, et al. Camonsertib in DNA damage response-deficient advanced solid tumours: phase 1 trial results. Nat Med 2023;29:1400–11.

[16] Saxena S, Zou L. Hallmarks of DNA replication stress. Mol Cell 2022;82:2298–314. 10.1016/j.molcel.2022.05.004.

[17] Toledo LI, Altmeyer M, Rask MB, et al. XATR prohibits replication catastrophe by preventing global exhaustion of RPA. Cell 2013;155:1088. 10.1016/j.cell.2013.10.043.

[18] Ngoi NYL, Pilié PG, McGrail DJ, et al. Targeting ATR in patients with cancer. Nat Rev Clin Oncol 2024;21:278–93. 10.1038/s41571-024-00863-5.

[19] Ng CKY, Dazert E, Boldanova T, et al. Integrative proteogenomic characterisation of hepatocellular carcinoma across etiologies and stages. Nat Commun 2022;13:2436. 10.1038/s41467-022-29960-8.

[20] Dreyer SB, Upstill-Goddard R, Paulus-Hock V, et al. Targeting DNA damage response and replication stress in pancreatic cancer. Gastroenterology 2021;160:362–77.

[21] Tian T, Bu M, Chen X, et al. The ZATT-TOP2A-PICH axis drives extensive replication fork reversal to promote genome stability. Mol Cell 2021;81:198–211.

[22] Selemenakis P, Sharma N, Uhrig ME, et al. RAD51AP1 and RAD54L can underpin two distinct RAD51-dependent routes of DNA damage repair via homologous recombination. Front Cell Dev Biol 2022;10:866601.

[23] Broutier L, Mastrogiovanni G, Verstegen MMA, et al. Human primary liver cancer – derived organoid cultures for disease modelling and drug screening. Nature Publishing Group 2017;23. 10.1038/nm.4438.

[24] Nuciforo S, Fofana I, Matter MS, et al. Organoid Models of Human Liver Cancers Derived from Tumour Needle Biopsies. Cell Rep 2018;24:1363–76. 10.1016/j.celrep.2018.07.001.

[25] Zou Z, Lin Z, Wu C, et al. Micro-Engineered Organoid-on-a-Chip Based on Mesenchymal Stromal Cells to Predict Immunotherapy Responses of HCC Patients. Advanced Science 2023;10:1–12. 10.1002/advs.202302640.

[26] Van Hemelryk A, Mout L, Erkens-Schulze S, et al. Modelling prostate cancer treatment responses in the organoid era: 3D environment impacts drug testing. Biomolecules 2021;11:1572.

[27] Steinmann SM, Lazzari M, Kleinle A, et al. The Novel HSF1 Inhibitor NXP800 Exhibits Robust Antitumor Activity in Hepatocellular Carcinoma. Int J Mol Sci 2026;27. 10.3390/ijms27062781.

[28] Takahashi N, Kim S, Schultz CW, et al. Replication Stress Defines Distinct Molecular Subtypes Across Cancers. Cancer Research Communications 2022;2:503–17. 10.1158/2767-9764.crc-22-0168.

[29] Woodward AM, Goehler T, Luciani MG, et al. Excess Mcm2-7 licenses dormant origins of replication that can be used under conditions of replicative stress. J Cell Biol 2006;173:673–83.

[30] Buisson R, Boisvert JL, Benes CH, et al. Distinct but concerted roles of ATR, DNA-PK, and Chk1 in countering replication stress during S phase. Mol Cell 2015;59:1011–24.

[31] Young LA, O’Connor LO, de Renty C, et al. Differential activity of ATR and WEE1 inhibitors in a highly sensitive subpopulation of DLBCL linked to replication stress. Cancer Res 2019;79:3762–75.

[32] Mahdi H, Hafez N, Doroshow D, et al. Ceralasertib-mediated ATR inhibition combined with olaparib in advanced cancers harbouring DNA damage response and repair alterations (olaparib combinations). JCO Precis Oncol 5: PO. 20.00439 2021.

[33] Reichert ZR, Devitt ME, Alumkal JJ, et al. Targeting resistant prostate cancer, with or without DNA repair defects, using the combination of ceralasertib (ATR inhibitor) and olaparib (the TRAP trial). 2022.

[34] Simpkins F, Nasioudis D, Wethington SL, et al. Combination ATR and PARP Inhibitor (CAPRI): A phase 2 study of ceralasertib plus olaparib in patients with recurrent, platinum-sensitive epithelial ovarian cancer (cohort A). 2024.

[35] Elayapillai SP, Dogra S, Lausen J, et al. ATR inhibition increases reliance on PARP-mediated DNA repair revealing an improved therapeutic strategy for cervical cancer. Gynecol Oncol 2024;191:182–93.

[36] Thangaretnam K, Islam MO, Lv J, et al. WEE1 inhibition in cancer therapy: Mechanisms, synergies, preclinical insights, and clinical trials. Crit Rev Oncol Hematol 2025;211. 10.1016/j.critrevonc.2025.104710.

[37] Bukhari AB, Lewis CW, Pearce JJ, et al. Inhibiting Wee1 and ATR kinases produces tumour-selective synthetic lethality and suppresses metastasis. Journal of Clinical Investigation 2019;129:1329–44. 10.1172/JCI122622.

[38] Giotti B, Chen SH, Barnett MW, et al. Assembly of a parts list of the human mitotic cell cycle machinery. J Mol Cell Biol 2019;11:703–18. 10.1093/jmcb/mjy063.

[39] Guo Y, Wang J, Benedict B, et al. Targeting CDC7 potentiates ATR-CHK1 signalling inhibition through induction of DNA replication stress in liver cancer. Genome Med 2021;13:1–15.

[40] Jia X, Zhu X, Chen S, et al. Comprehensive multi-omics analyses expose a precision therapy strategy that targets replication stress in hepatocellular carcinoma using WEE1 inhibition. J Adv Res 2025.

[41] Donne R, Saroul-Ainama M, Cordier P, et al. Replication stress triggered by nucleotide pool imbalance drives DNA damage and cGAS-STING pathway activation in NAFLD. Dev Cell 2022;57:1728–41.

[42] Griffiths CD, Zhang B, Tywonek K, et al. Toxicity profiles of systemic therapies for advanced hepatocellular carcinoma: a systematic review and meta-analysis. JAMA Netw Open 2022;5:e2222721–e2222721.

[43] Böhly N, Schmidt AK, Zhang X, et al. Increased replication origin firing links replication stress to whole chromosomal instability in human cancer. Cell Rep 2022;41. 10.1016/j.celrep.2022.111836.

[44] Merry E, Gourley C. Targeting DNA Damage Repair Pathways Beyond PARP Inhibition. Target Oncol 2025;20:937–53. 10.1007/s11523-025-01183-z.

[45] Ruiz S, Mayor-Ruiz C, Lafarga V, et al. A Genome-wide CRISPR Screen Identifies CDC25A as a Determinant of Sensitivity to ATR Inhibitors. Mol Cell 2016;62:307–13. 10.1016/j.molcel.2016.03.006.

[46] Yang X, Wang H, Zhang L, et al. Novel roles of karyopherin subunit alpha 2 in hepatocellular carcinoma. Biomedicine and Pharmacotherapy 2023;163. 10.1016/j.biopha.2023.114792.

[47] Jiang P, Tang Y, He L, et al. Aberrant expression of nuclear KPNA2 is correlated with early recurrence and poor prognosis in patients with small hepatocellular carcinoma after hepatectomy. Medical Oncology 2014;31. 10.1007/s12032-014-0131-4.

[48] Ding K, Liu L, Yong W, et al. Bioinformatics analysis and experimental studies reveal KPNA2 as a novel biomarker of hepatocellular carcinoma progression and telomere maintenance. Eur J Med Res 2025;30. 10.1186/s40001-025-02866-z.

[49] Taniguchi H, Chakraborty S, Takahashi N, et al. ATR inhibition activates cancer cell cGAS/STING-interferon signalling and promotes antitumor immunity in small-cell lung cancer. vol. 10. 2024.

