## Supplementary material for "High regeneration-associated stress defines a distinct HCC subgroup with therapeutically exploitable vulnerabilities": Methods, Supplementary Figures and Table

### Supplemental material and methods

#### Material and methods

##### *Human tissue*

Liver tumour samples were obtained from patients undergoing surgical resection (1-3 $cm^2$) or liver biopsy (1 mm x 4-10 mm) at the University Hospital Zurich, Zurich, Switzerland and IRCCS Humanitas Research Hospital, Rozzano, Milan, Italy. Written informed consent was obtained from all patients. The study was conducted under the following ethical approval (KEK-ZH-2013-0382, KEK-ZH-2014-03-20) in Switzerland and Italy (no. 4174). Patient samples were placed in advanced DMEM/F12 (Gibco,

12634) for transport and processed to develop organoids within 10-30 min after reception of the tissue. One-half of the specimen was embedded in O.C.T. (Tissue-Tek) and frozen according to standard procedures, while the other half was further processed for organoid generation.

*Organoid culture*

Tumour tissue was first minced, followed by removal of red blood cells by red blood cell lysis buffer (Biosciences, 786-649) for 5 min at RT and then washed by a 3 min centrifugation step (300 *x g,* 3 min). Tissue was digested in 2.5 mg/ml collagenase type IV (Sigma-Aldrich, C5138), 0.1 mg/ml DNase I (Roche, 10104159001), and advanced DMEM/F12 (Gibco, 12634) at 37°C for 20-70 min. Digestion was stopped until the presence of small cell clusters by the addition of advanced DMEM/F12 (Gibco) and by centrifugation (300 *x g*, 5 min). Digestion time varied between patient samples depending on size, fibrotic tissue and viability of the tumour sample. The cells were then mixed with the culture medium and plated in ultra-low attachment plates (Corning Costar, 3473). Culture medium was composed of advanced DMEM/F12 (Gibco, 12634), 100 µg/ml Primocin (InvivoGen, ant-pm-05), 10 mM Hepes (Gibco, 15630-056), B27 supplement (Gibco, 17504044), 10 mM Nicotonamide (Sigma-Aldrich, N0636), 1.25 mM N-Acetyl-L-Cysteine (Sigma-Aldrich, A9165), 1 nM Gastrin I (MedChemExpress, HY-P1097), 10 $\mu$M SB202190 (Selleckchem, S1077), 50 ng/ml EGF (Peprotech, AF-100-15), 1 ng/ml FGF-basic (Peprotech, AF-100-18B), 20 ng/ml FGF-10 (Peprotech, 100-26-100), 25 ng/ml HGF (Peprotech, 100-39H), 10 µM Y-27632 (Selleckchem, S1049), 0.5 µM A83-01 (Sigma-Aldrich, SML0788), 1 $\mu$M PGE2 (Selleckchem, S3003), 10 µM Forskolin (MedChemExpress, HY-15371) and 10 % Wnt3a/R-Spondin1/Noggin conditioned medium (*in-house*). Organoids were passaged by dissociation by incubation in TryplE (Gibco, 12563) for 15 min at 37°C. Dissociation was stopped by adding advanced DMEM/F12 (Gibco) and by centrifugation (300 *x g*, 5 min). The first passage was done 7-14 days after initial plating. Partial medium change was done every 2-3 days by removing 1/4 of the medium and replacing it with fresh medium. Organoids were frozen at regular passages, once the fourth passage was reached, after dissociation in 1x TryplE™ select (Gibco, 12563) and resuspended in Recovery Cell Culture Freezing Medium (Gibco, 12648-010) before freezing. All organoid lines were kept in culture at least until the 20$^{th}$ passage. Organoid lines P151, P373, P558 and P636 were given by IRCCS Humanitas Research Hospital, Rozzano, Italy, established as previously published and adapted to the culture medium mentioned above [1]. All organoid lines were incubated at 37°C in a humidified atmosphere with 5% $CO_2$ and tested for Mycoplasma (Promokine, pk-CA91-1024).

*HepaRG cell culture and 3D spheroid formation*

The human hepatic non-differentiated HepaRG cell line (Biopredic, MTA_HRG_V3_2018_07012028) was maintained in William's E Medium with GlutaMax (Gibco, 32551020) supplemented with 1x HepaRG growth medium supplements with antibiotics (Biopredic, ADD710C). Cells were passaged before confluency using TryplE™ select (Gibco, 12563). HepaRG spheroids were developed by the droplet technique, with 500 cells/drop. Drops of 20 µl were pipetted on an inverted lid from a 10 cm petri dish (Falcon, 351029). Drops were then hung by inverting the lid and incubated for 2 days, before transferring the spheroids to an ultra-low attachment plate (Corning, 3473). The cells were incubated at 37°C in a humidified atmosphere with 5% $CO_2$ and tested for Mycoplasma (Promokine, pk-CA91-1024).

*Weighted gene co-expression network analysis*

The published bulk RNA-seq dataset from Ng. *et al.* was used as a training dataset [2], while the publicly available dataset from TCGA (LIHC) served as a validation dataset. Access to the first dataset was provided by EGA and the corresponding author. The weighted gene co-expression network analysis (WGCNA) was performed using the WGCNA *R* package (version 1.73) [3]. Briefly, the WGCNA involves several key steps: first, correlations are calculated between each pair of genes, and strong correlations are then organised into a connected network. To delimit modules within this newly defined network, the dissimilarity between genes is computed based on the topological overlap measure (TOM). This measure quantifies the number of shared neighbours for each pair of genes, which is then normalised. A high TOM indicates that the two genes exhibit similar expression patterns, which is subsequently converted into a dissimilarity score (1-TOM). Genes are then clustered based on their TOM, and the clusters are divided into modules using hierarchical clustering. The final step of the WGCNA involves merging highly similar modules by computing module eigengenes, which represent the first principal component of gene expression of each module, effectively serving as a one-dimensional vector. A clustering can be applied on the newly defined modules and merge highly correlated modules to avoid biological repetitions or redundancies within modules.

For the implementation of the WGCNA, gene filtering was performed based on a differential expression analysis between healthy donors and HCC tissue, resulting in 2539 genes. TPM-normalised RNA-seq served as input data for the WGCNA. A soft-thresholding power was chosen to approximate scale-free topology. An adjacency matrix was then computed and transformed into a topological overlap matrix (TOM) to cluster genes into modules via average linkage hierarchical clustering. Modules were defined through a dynamic tree-cut algorithm, and module eigengenes were correlated with clinical traits and GO terms to identify relevant modules. Gene significance and module membership were evaluated to prioritise the module, which was relevant to replication stress.

*RNA extraction and bulk RNA-sequencing*

Total RNA from tumour organoids was extracted using the RNAeasy Micro Kit (Qiagen, 74004) following the manufacturer's instructions. And the total RNA from biopsies and surgical resections was extracted using the MaxWell RSC SimplyRNA tissue kit (Promega, AS1340) with the MaxWell RSC Instrument (Promega, AS4500) according to the manufacturer's instructions. Quantification of the extracted RNA was performed using the Qubit Fluorometer (ThermoFisher Scientific). 200 ng of total RNA was used for RNA-seq library prep with the SMARTer Stranded Titak RNA-seq kit (Takara Bio), and sequencing was performed on an Illumina NovaSeq X Plus with paired-end 150 bp at the Functional Genomics Centre Zurich following the manufacturer's instructions.

*Mid-throughput drug screening*

Mid-throughput drug screen was performed using the CyBio Felix liquid handler (Analytik Jena). Organoids were first dissociated into a single cell suspension in Cell Detachment Solution (Accutase, AT-104) for 20 min at 37°C. 500 cells/well were then seeded in PrimeSurface U-bottom, ultra-low attachment 384 well-plates (SBio, MS-9384UZ) in adapted culture medium. Adapted culture medium was composed of the culture medium but inhibitors which might interfere the drug response were removed, *i.e.*, advanced DMEM/F12 (Gibco, 12634), 100 µg/ml Primocin (InvivoGen, ant-pm-05), 10 mM Hepes (Gibco, 15630-056), B27 supplement (Gibco, 17504044), 10 mM Nicotonamide (Sigma-Aldrich, N0636), 1.25 mM N-Acetyl-L-Cysteine (Sigma-Aldrich, A9165), 1 nM Gastrin I (MedChemExpress, HY-P1097), 50 ng/ml EGF (Peprotech, AF-100-15), 1 ng/ml FGF-basic (Peprotech, AF-100-18B), 20 ng/ml FGF-10 (Peprotech, 100-26-100), 25 ng/ml HGF (Peprotech, 100-39H), 1 µM PGE2 (Selleckchem, S3003), 10 µM Forskolin (MedChemExpress, HY-15371) and 10 % Wnt3a/R-Spondin1/Noggin conditioned medium (*in-house*). Organoids were treated with a customised drug library including 68 targeted or chemotherapeutic agents (MedChemExpress, Sup. Table 1) in 5 doses of serial dilution (1 nM to 10 µM) in technical quadruplicates, 2 and 7 days after cell seeding, in drug screen medium. Drug screen medium contained *in vitro* nutrition medium (Cell Culture Technologies, VNU-0.5), 10 % Fetal Bovine Serum Albumin (Gibco, 2575623) and 100 µg/ml Primocin (InvivoGen, ant-pm-05). Cell viability readout was performed using Cell Titer Glo® Luminescent Cell Viability assay (Promega, G7572), 10 days after the first drugging step. Measurement of the luminescence was performed on a multiplate reader (Tecan Infinite 200 Pro). Controls included organoids treated with 0.1 % DMSO as a negative control, *i.e.*, 100 % cell viability, and organoids treated with 1 µM staurosporine as a positive control for cell death, *i.e.*, 0 % viability. All graphs and statistical analyses were performed using GraphPad Prism (version 10.4.2) software.

*Drug screening validation*

The validation of the drug screen experiments was done with the same pipeline as the "Mid-throughput drug screening" mentioned above, but including all cell models together and with the following modifications. Cell seeding was performed on roundbottom, ultra-low attachment 96-well plates (Corning, 7007). Technical triplicates were seeded on the plate. The following compounds were tested: camonsertib (MedChemExpress, HY-139609), ceralasertib (Selleckchem, S7693), elimusertib (MedChemExpress, HY-101566), berzosertib (Selleckchem, S7102), SAR-020106 (Selleckchem, S7740), rabusertib (MedChemExpress, HY-14720), azenosertib (MedChemExpress, HY-132295), olaparib (MedChemExpress, HY-10162), niraparib (MedChemExpress, HY-10619), lartesertib (Selleckchem, E1057), AZD1390 (MedChemExpress, HY-109566), sorafenib (MedChemExpress, HY-10201), lenvatinib (MedChemExpress, HY-10981), pemetrexed (Selleckchem, S5971), methotrexate (MedChemExpress, HY-14519), staurosporine (MedChemExpress, HY-15141). A non-linear regression was computed based on the 5-point dose response (1 nM to 10 μM), and % normalised viability and quality were evaluated based on $R^2$. $IC_{50}$ and AUC were calculated from the non-linear regression. The average AUC of the technical triplicates was used for statistical analysis. All graphs and statistical analyses were performed using GraphPad Prism (10.4.2) software or *R* (4.4.2) with the pheatmap *R* package (1.0.12) [4,5].

*Combination drug testing*

Combination drug testing was performed with the same pipeline as the "Drug screening validation" mentioned above, but with the following modifications. Cell viability was measured by luminescence using Cell Titer Glo® Luminescent Cell Viability assay (Promega, G7572), 10 days after the first drugging step. Drug response graphs were performed using GraphPad Prism (version 10.4.2) software, and drug synergistic scores were calculated using the SynergyFinder *R* package (3.14.0).

*DNA fibre spreading assay*

Organoids and spheroids were pulse labelled with 30 $\mu$M of the thymidine analogue, 5-chloro-2'-deoxyuridine (CldU; Sigma-Aldrich, C6891) for 35 min, gently washed with warm PBS, followed by incubation with 250 µM of 5'-iodo-2'-deoxyuridine (IdU; Sigma-Aldrich, I7125) for 30 min at 37°C. Ongoing replication was stopped by washing 3 times with cold PBS. Single cell suspension was obtained by incubating the organoids and spheroids in TrypLE$^{TM}$ Select for 20 min at 37°C. Cells were then resuspended in cold PBS at 3.5 x $10^5$ cells/ml. Cells were lysed by mixing 3 µl of the cell suspension with 7 µl of lysis buffer (200 nM Tris HCl, pH 7.5, 50 mM EDTA, and 0.5% [w/vol] SDS) on a microscope slide (Epredia, J1800AMNZ). After 6 min of incubation at RT, the slides were tilted at a 30-45° angle to stretch the DNA fibres onto the slide. The DNA spreads were then air-dried, fixed in 3:1 methanol/acetic acid, and stored at 4°C overnight. The DNA spreads were incubated in 2.5 M HCl for 1 h at RT to denature the DNA and then washed 5 times with PBS and blocked with 2% (Merck, 05470) in 0.2 % PBST (PBS and Tween-20) for 40 min at RT. CldU and IdU were stained with rat anti-BrdU/CldU antibodies (Abcam, ab6326, RRID: AB-305426, 1:400) and mouse anti-BrdU/IdU antibodies (BD Biosciences, 347580, RRID: AB-400326, 1:80), respectively, for 2.5 h at RT. After washing 5 times with 0.2 % PBST (PBS and Tween-20), the slides were incubated with the secondary antibodies, goat anti-rat Alexa555 (Thermo Fisher Scientific, A-21434, RRID: AB-2535855, 1:200) and goat anti-mouse Alexa488 (Thermo Fisher Scientific, A-11001, RRID: AB-2534069, 1:300) for 2 h at RT in the dark. After washing 5 times with 0.2 % PBST (PBS and Tween-20), the slides were mounted by the addition of 30 µL Prolong$^{TM}$ Gold antifade mounting medium (Thermo Fischer Scientific, P36930). The distribution of the fibres' length is shown by one replicate (n = 150 structures/condition per replicate), while the medians of three independent replicates are shown (black dots) with the corresponding average (black line).

The fibres were imaged using a Leica DM6 microscope (HCX PL APO, 63x objective). Total fibre track length and sister fork length were measured using the line tool in Fiji/ImageJ (2.1.0/1.53c). 150 molecules were measured by technical replicate. Origin firing was assessed using the cell counter plugin in Fiji/ImageJ (2.1.0/1.53c). At least 500 events were counted per technical replicate. Researchers were blinded with respect to the treatment during image analysis. Statistical analysis was performed on the median of the technical triplicates using GraphPad Prism (10.4.2) software.

*Automated immunostaining*

Immunohistochemistry was performed on FFPE sections from embedded cell pellets following the standard procedures established at the Department of Pathology and Molecular Pathology. Briefly, FFPE sections were rehydrated, followed by a heat-induced antigen retrieval step in either Tris/EDTA/BORAT or citrate buffer. Incubation in Ventana buffer and staining were performed on a Benchmark Ultra immunohistochemistry robot (Ventana Instruments) using Optiview DAB Detection Kits (Ventana) or on a Bond MAX (Leica, Wetzlar, Germany). Primary antibodies: anti-cleaved caspase 3 (Cell Signaling Technology, 9664, RRID: AB_2070042, 1:100); anti-KPNA2 (Abcam, ab289858, EPR25248-95, 1:5000); anti-cytokeratin 7 (Abcam, ab183344, RRID: AB_2936915, 1:100); anti-hepatocyte specific antigen (Vector Laboratories, VP-H907, RRID: RRID:AB_2336504, 1:100). Image quantification of Cl. Casp. 3 staining was done using QuPath (v0.5.0) [6]. Semi-quantification of KPNA2 staining was done by one blinded researcher evaluating the nuclear intensity (negative: 0, weak: 1, moderate: 2, strong: 3) and the proportion of KPNA2 positive cells (0%: 0, <1%: 1, [1-10%]: 2; [10-33%]: 3) in duplicates for the tissue microarray and at least 8 organoids for the HCC-Org analysis (KPNA2 IHC score).

*Immunofluorescence staining and mitotic figures analysis*

Cells were treated for 72 h with 500 nM Ceralasertib or 0.1 % DMSO as a negative control. HCC-Org were washed in PBS, fixed for 40 min at RT with 4 % paraformaldehyde and washed in PBS. The organoids were dehydrated in 100 % ethanol and embedded in paraffin prior to sectioning. Three µm sections were dewaxed in xylene and rehydrated, followed by a heat-induced antigen retrieval step in citrate buffer. Heat-induced antigen retrieval was performed using 10 mM sodium citrate buffer (Sigma-Aldrich, 71402), 0.05 % tween 20, pH 6, for 10 min. The cuts were then permeabilised and blocked in blocking solution: 0.2 % Triton X-100 (Sigma-Aldrich, 93443), 5 % bovine serum albumin (Sigma-Aldrich, A3912) in PBS for 1 h at RT. Next, sarcospheres were immunostained with rabbit anti-beta-tubulin (clone D3U1W; Cell Signaling Technology, 86298, RRID: AB_2715541). After performing three washes in PBS, HCC-Org were immunostained with the secondary antibodies: Alexa FluorTM 488 goat anti-mouse IgG (Thermo Fischer Scientific, A11001, RRID: AB_2534069) and 1ug/ml 4′,6-diamidin-2-phenylindol (DAPI; Sigma-Aldrich, D9542). Slides were air-dried, mounted with ProLongTM Gold Antifade mounting medium (Invitrogen, P36930) and covered with a coverslip. Imaging was performed at the Leica Stellaris 5, using a 63x magnification and analysed in Fiji/ImageJ (v1.54f). Researchers were blinded with respect to the treatment during imaging and image analysis. Unblinding was performed immediately before final data analysis and interpretation. Abnormal mitoses were defined by atypical spindle symmetry, asymmetrical relative proportion or polarity, metaphase plate asymmetry, multipolar mitosis, or abnormal chromosomal segregation (bridging, lagging or distribution symmetry). The proportion of abnormal mitotic figures was calculated as a percentage across three technical replicates (n ≥ = 25 events/condition per replicate).

*Statistical analysis*

Statistical analysis was performed as mentioned in the figure legends, using either GraphPad Prism (version 10.4.2) software or *R* (4.4.2) [5].

*Data availability*

Quantitative data, including all individual data points shown in the graph, are provided as a supporting data values file. *In-house* bulk RNA-seq data will be available upon request.

Supplemental figures

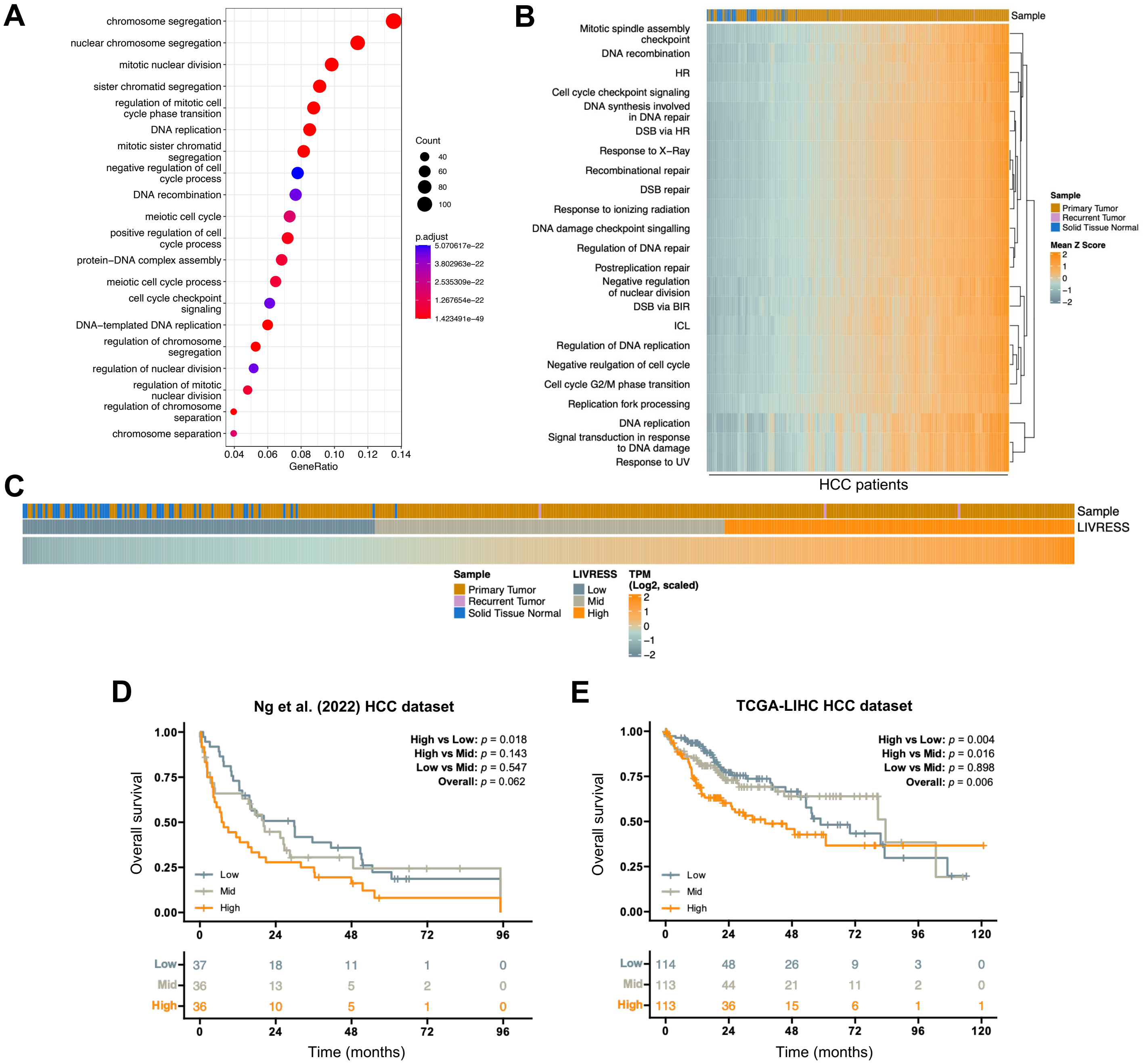
A
chromosome segregation
nuclear chromosome segregation
mitotic nuclear division
sister chromatid segregation
regulation of mitotic cell cycle phase transition
DNA replication
mitotic sister chromatid segregation
negative regulation of cell cycle process
DNA recombination
meiotic cell cycle
positive regulation of cell cycle process
protein−DNA complex assembly
meiotic cell cycle process
cell cycle checkpoint signaling
DNA−templated DNA replication
regulation of chromosome segregation
regulation of nuclear division
regulation of mitotic nuclear division
regulation of chromosome separation
chromosome separation
0.04 0.06 0.08 0.10 0.12 0.14
GeneRatio
Count
40
60
80
100
p.adjust
5.070617e−22
3.802963e−22
2.535309e−22
1.267654e−22
1.423491e−49
B
Mitotic spindle assembly checkpoint
DNA recombination
HR
Cell cycle checkpoint signaling
DNA synthesis involved in DNA repair
DSB via HR
Response to X-Ray
Recombinational repair
DSB repair
Response to ionizing radiation
DNA damage checkpoint singalling
Regulation of DNA repair
Postreplication repair
Negative regulation of nuclear division
DSB via BIR
ICL
Regulation of DNA replication
Negative reulgation of cell cycle
Cell cycle G2/M phase transition
Replication fork processing
DNA replication
Signal transduction in response to DNA damage
Response to UV
HCC patients
Sample
Sample
Primary Tumor
Recurrent Tumor
Solid Tissue Normal
Mean Z Score
2
1
0
−1
−2
C
Sample
LIVRESS
Sample
Primary Tumor
Recurrent Tumor
Solid Tissue Normal
LIVRESS
Low
Mid
High
TPM (Log2, scaled)
2
1
0
−1
−2
D
Ng et al. (2022) HCC dataset
High vs Low: p = 0.018
High vs Mid: p = 0.143
Low vs Mid: p = 0.547
Overall: p = 0.062
Overall survival
1.00
0.75
0.50
0.25
0.00
Low
Mid
High
0 24 48 72 96
Low 37 18 11 1 0
Mid 36 13 5 2 0
High 36 10 5 1 0
0 24 48 72 96
Time (months)
E
TCGA-LIHC HCC dataset
High vs Low: p = 0.004
High vs Mid: p = 0.016
Low vs Mid: p = 0.898
Overall: p = 0.006
Overall survival
1.00
0.75
0.50
0.25
0.00
Low
Mid
High
0 24 48 72 96 120
Low 114 48 26 9 3 0
Mid 113 44 21 11 2 0
High 113 36 15 6 1 1
0 24 48 72 96 120
Time (months)

**Fig. S1**

**Fig. S1. HCC patients with higher regeneration stress-associated have more aggressive disease and poorer survival.** (A) Pathway enrichment analysis of the module identified by weighted gene co-expression analysis (WGCNA). (B) Heatmap with clustering using GO terms (TCGA LIHC dataset). (C) Ranking of HCC patients based on the LIVRESS score. LIVRESS score is calculated using the mean Z score (transcripts per million (TPM)). (D,E) Kaplan Meier survival curves with patient number from Ng *et al*. dataset (D) [2] and from the TCGA-LIHC dataset (E). Log-rank test for survival. P-value indicated.

A

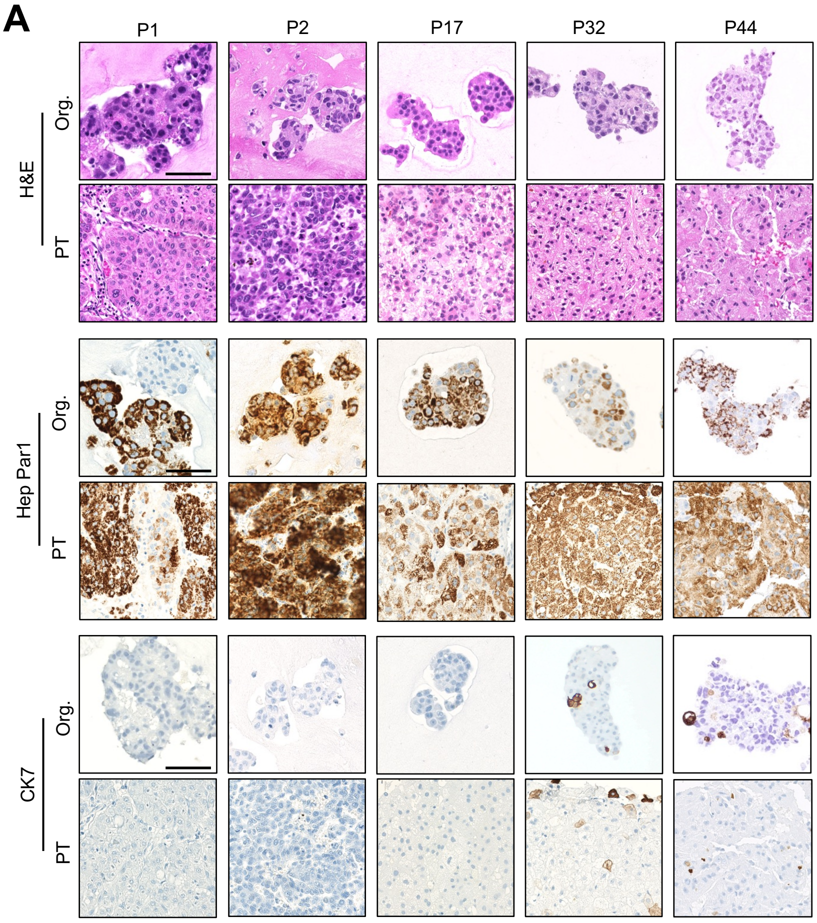

B

| Patient ID | Growth pattern (PT) | Growth pattern (Org) | WHO grade (PT) | WHO grade (Org) |
|---|---|---|---|---|
| P1 | solid/pd/trabecular | solid/pd | G2 | G2 |
| P2 | solid | solid | G3 | G3 |
| P17 | solid | solid/pd | G2 | G2 |
| P32 | solid/clear cell | solid | G2 | G2 |
| P44 | solid/pd/trabecular | solid/steatohepatitic | G2 | G2/G3 |

Org, HCC-organoids; pd, pseudoglandular; PT, parental tumour

**Fig. S2**

**Fig. S2. HCC organoids (HCC-Org) recapitulate the heterogeneous regeneration stress landscape of clinical cohorts**. (A) Morphological and histological characterisation of the HCC-Org and parental tumour (PT) using immunohistochemical HCC and biliary markers. Scale bar, 100 μm. (B) Table of pathological characterisation between the PT and the HCC-Org, including growth pattern and WHO grade. The pathologist blindly performed the assessment.

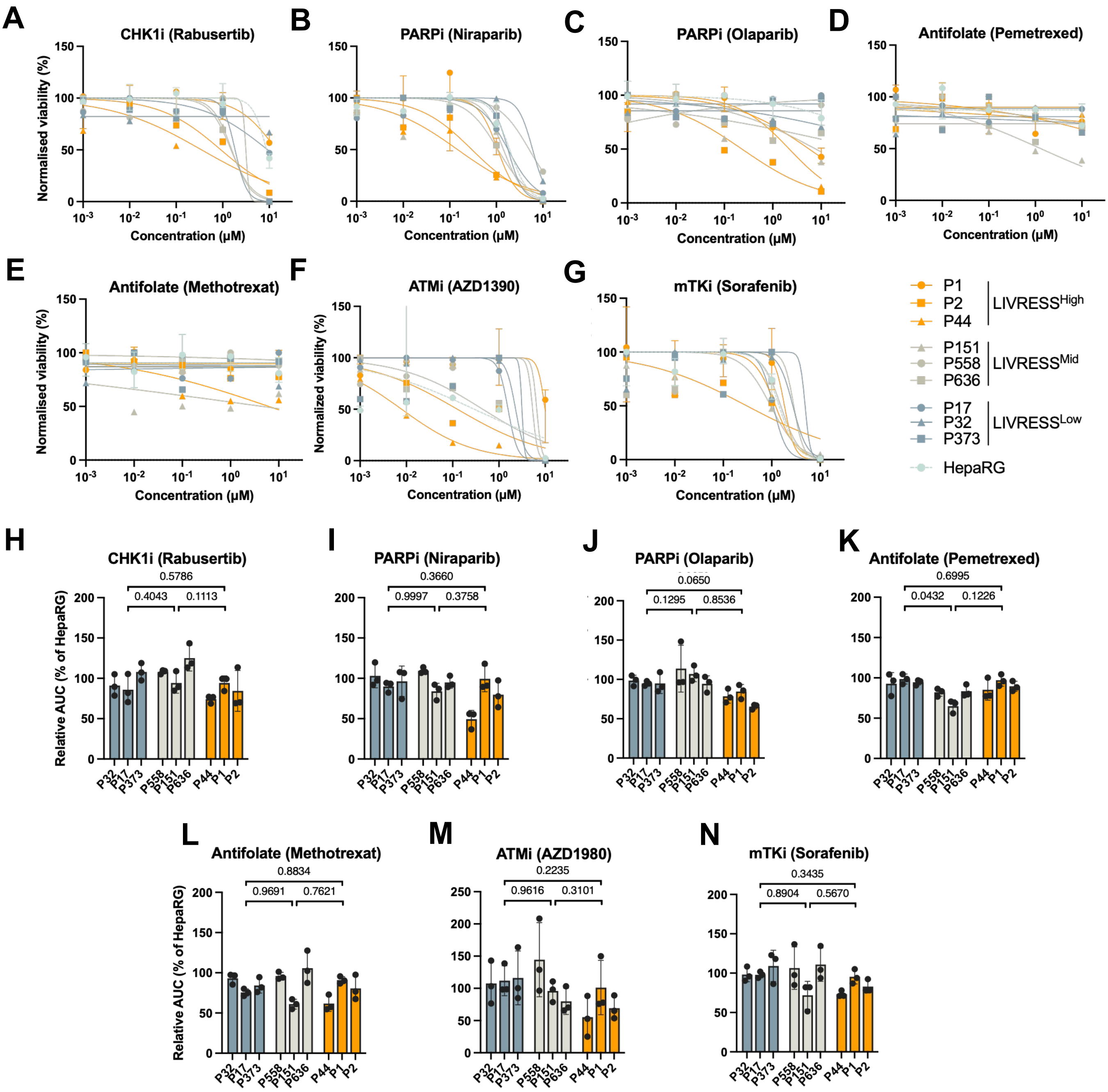

A
CHK1i (Rabusertib)
B
PARPi (Niraparib)
C
PARPi (Olaparib)
D
Antifolate (Pemetrexed)
E
Antifolate (Methotrexat)
F
ATMi (AZD1390)
G
mTKi (Sorafenib)
Normalised viability (%)
Normalized viability (%)
Concentration (µM)
P1
P2
P44
LIVRESS^High
P151
P558
P636
LIVRESS^Mid
P17
P32
P373
LIVRESS^Low
HepaRG
H
CHK1i (Rabusertib)
0.5786
0.4043
0.1113
I
PARPi (Niraparib)
0.3660
0.9997
0.3758
J
PARPi (Olaparib)
0.0650
0.1295
0.8536
K
Antifolate (Pemetrexed)
0.6995
0.0432
0.1226
L
Antifolate (Methotrexat)
0.8834
0.9691
0.7621
M
ATMi (AZD1980)
0.2235
0.9616
0.3101
N
mTKi (Sorafenib)
0.3435
0.8904
0.5670
Relative AUC (% of HepaRG)
P32
P17
P373
P558
P151
P636
P44
P1
P2

**Fig. S3**

**Fig. S3. Functional screening identifies ATR as a vulnerability in high LIVRESS HCC-Org.** (A-N) Viability dose-response curves (A-G) and relative AUC (H-N, normalised to HepaRG) from drug validation experiments in HCC-Org and HepaRG spheroids (normalised to DMSO control). Statistics: nested one-way ANOVA with Tukey's multiple comparisons test. Average and SD from three independent replicates are shown. *P*-values indicated.

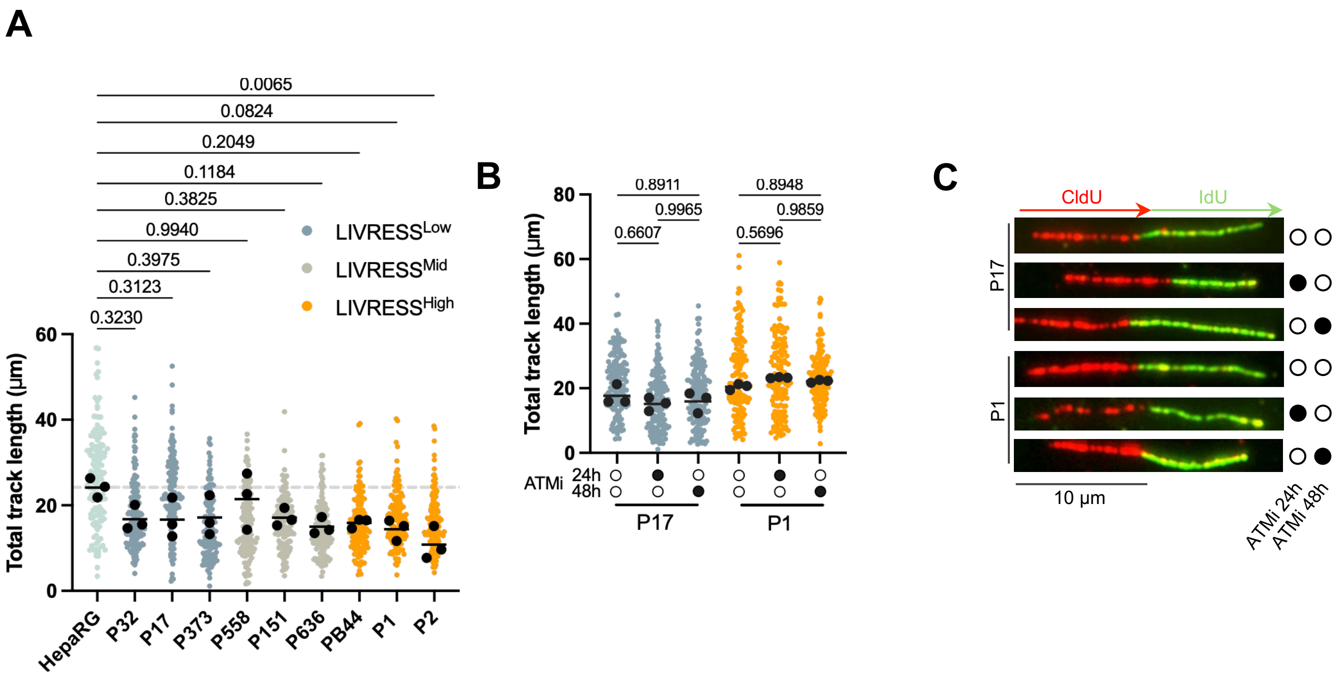

A
0.0065
0.0824
0.2049
0.1184
0.3825
0.9940
0.3975
0.3123
0.3230
LIVRESS^Low
LIVRESS^Mid
LIVRESS^High
Total track length (µm)
HepaRG
P32
P17
P373
P558
P151
P636
PB44
P1
P2
B
Total track length (µm)
0.8911
0.9965
0.6607
0.8948
0.9859
0.5696
ATMi
24h
48h
P17
P1
C
CldU
IdU
P17
P1
10 µm
ATMi 24h
ATMi 48h

Fig. S4

**Fig. S4. ATRi sensitivity is decoupled from replication fork dynamics and is linked to mitotic fragility.** (A) Baseline DNA fibre length in HCC-Org and HepaRG spheroids. (B, C) Total fibre length and representative images in P17/P2 ± ATMi (AZD1390, 100 nM). Scale, 10 µm. Data represent medians of 3 replicates (n=150 fibres/replicate). The distribution of the fibre length is always shown by one representative replicate, while the medians of three independent replicates are shown (black dots) with the corresponding average (black line). Statistics: One-way (A, C, D) or nested (B) ANOVA with Tukey's test. *P*-values indicated.

A
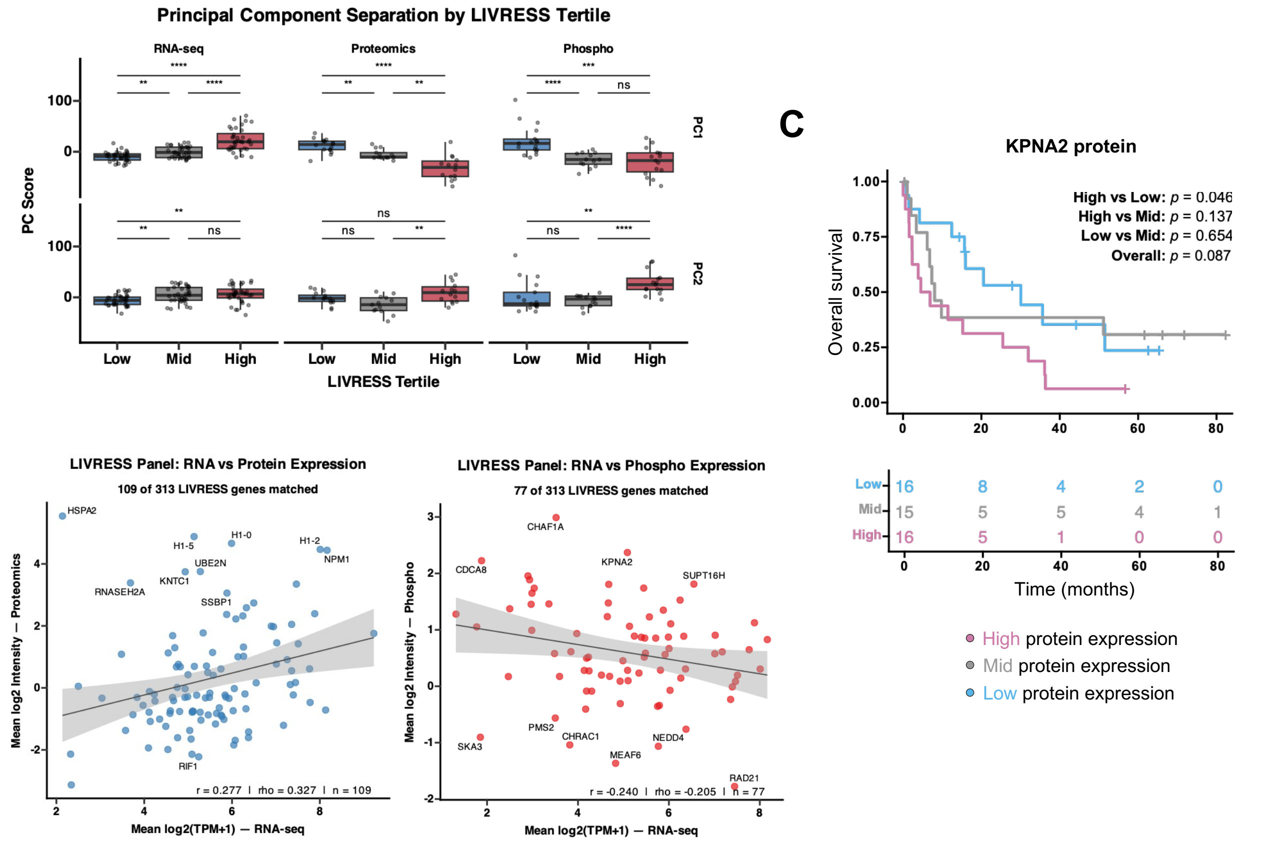

D

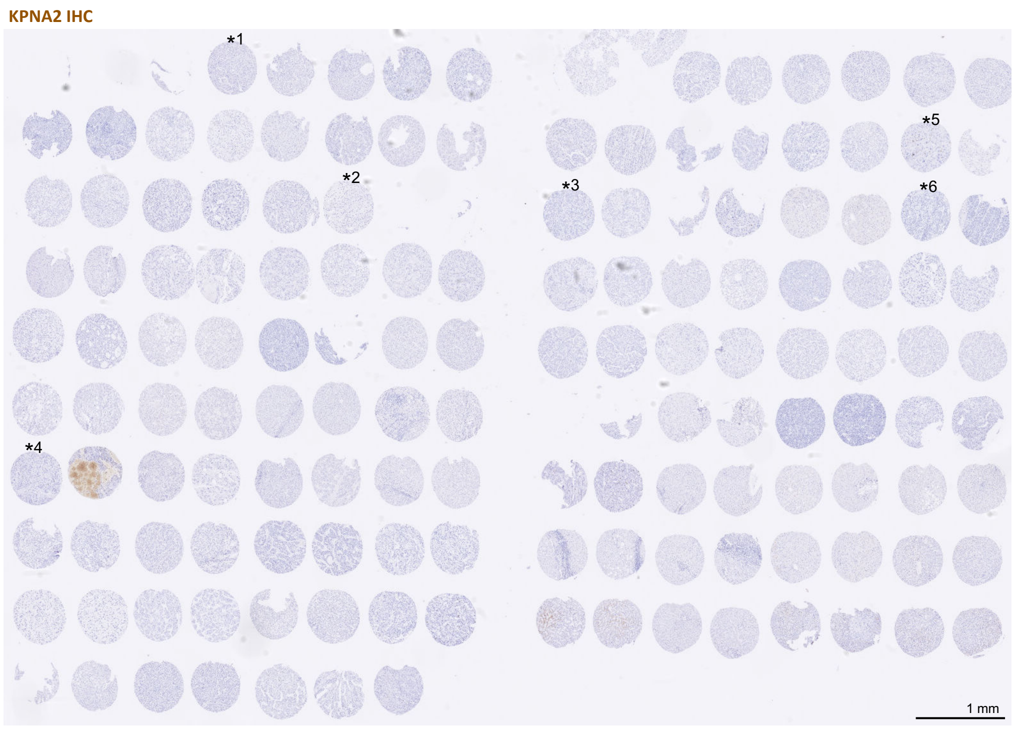

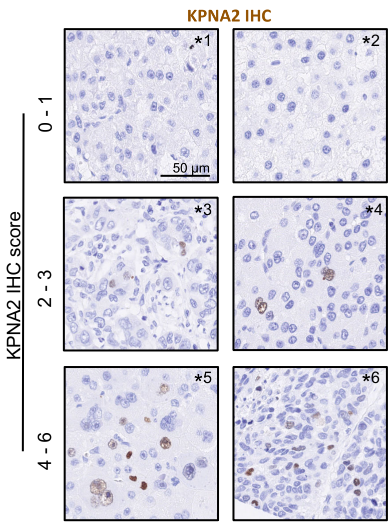

| KPNA2 IHC score | N° of patients |
|---|---|
| 0 - 1 | 36 |
| 2 - 3 | 15 |
| 4 - 6 | 7 |

**Fig. S5**

**Fig. S5. Multi-omic profiling identifies LIVRESS surrogate markers linked to poor HCC prognosis.** (A) boxplots of principal component analyses transcriptomic, proteomic and phosphoproteomic based on the LIVRESS score of HCC biopsies. (B) Scatter plot of correlation of LIVRESS genes (transcripts per million (TPM)) with matched LIVRESS proteins or with matched LIVRESS phosphorylation sites. (C) Kaplan Meier survival curves of KPNA2 protein expression with patient number. (D) IHC staining of KPNA2 on FFPE tissue microarray (TMA) and KPNA2 IHC score of HCC patients. *P*-value is indicated. Statistics: one-way ANOVA followed by Tukey's multiple comparisons test (A). Dataset [2].

Supplemental references

[1] Steinmann SM, Lazzari M, Kleinle A, et al. The Novel HSF1 Inhibitor NXP800 Exhibits Robust Antitumor Activity in Hepatocellular Carcinoma. Int J Mol Sci 2026;27. https://doi.org/10.3390/ijms27062781.

[2] Ng CKY, Dazert E, Boldanova T et al. Integrative proteogenomic characterisation of hepatocellular carcinoma across etiologies and stages. Nat Commun 2022;13:2436. https://doi.org/10.1038/s41467-022-29960-8.

[3] Langfelder P, Horvath S. WGCNA: an R package for weighted correlation network analysis. BMC Bioinformatics 2008:559.

[4] Kolde R. pheatmap: Pretty Heatmaps 2018.

[5] R Core Team R, others. R: A language and environment for statistical computing 2025.

[6] Bankhead P, Loughrey MB, Fernández JA, et al. QuPath: Open source software for digital pathology image analysis. Sci Rep 2017;7. https://doi.org/10.1038/s41598-017-17204-5.

[7] Nuciforo S, Fofana I, Matter MS, et al. Organoid Models of Human Liver Cancers Derived from Tumour Needle Biopsies. Cell Rep 2018;24:1363–76. https://doi.org/10.1016/j.celrep.2018.07.001.

Supplementary Table

*Table S1. Drug library*

| Position | Drug | Target | Position | Drug | Target |
|---|---|---|---|---|---|
| *A2* | Regorafenib | mTK inhibitor | *D4* | Cabozantinib | mTK inhibitor |
| *A3* | HA15 | HSPA5 inhibitor | *D5* | Elimusertib | ATR inhibitor |
| *A4* | GSK2606414 | PERK inhibitor | *D6* | Imatinib | mTK inhibitor |
| *A5* | AMG900 | AURKA inhibitor | *D7* | Rabusertib | CHK1 inhibitor |
| *A6* | Kira8 | IRE1α inhibitor | *D8* | Niraparib | PARP1/2 inhibitor |
| *A7* | Apoptozole | HSP70 inhibitor | *D9* | Pemetrexed | Antifolate |
| *A8* | Griseofulvin | Antifungal | *D10* | Irinotecan | Topoisomerase I inhibitor |
| *A9* | AZD1390 | ATM inhibitor | *D11* | Pazopanib | mTK inhibitor |
| *A10* | MKC3946 | IRE1α inhibitor | *E2* | Docetaxel | Taxane |
| *A11* | TD52 | CIP2A inhibitor | *E3* | Bortezomib | Proteasome inhibitor |
| *A12* | Oxaliplatin | Platinum | *E4* | Sorafenib | mTK inhibitor |
| *B2* | IBR2 | RAD51 inhibitor | *E5* | Vorinostat | HDAC inhibitor |
| *B3* | Gemcitabine | Antimetabolite | *E6* | Prexasertib | CHK1 inhibitor |
| *B4* | Amuvatinib | mTK, RAD51 inhibitor | *E7* | Tanespimycin | HSP90 inhibitor |
| *B5* | RP-6306 | PKMYT1 inhibitor | *E8* | Temozolomide | Alkylating agent |
| *B6* | VER-155008 | HSP70 inhibitor | *E9* | Chloroquine | Sulfonamide |
| *B7* | Onvansertib | PLK1 inhibitor | *E10* | Tauroursodeoxycholic acid | ER stress inhibitor |
| *B8* | SAR-020106 | CHK1 inhibitor | *F2* | Trabectedin | Alkylating agent |
| *B9* | LY294002 | PI3K/AKT inhibitor | *F3* | PUH71 | HSP90 inhibitor |
| *B10* | Rapamycin | MTOR inhibitor | *F4* | Novobiocin | Polymerase theta inhibitor |

| | | | | | |
|---|---|---|---|---|---|
| *B11* | GSK2656157 | PERK inhibitor | *F5* | Doxorubicin | Anthracyclin e |
| *B12* | Cisplatin | Platinum | *F6* | Lartesertib | ATM inhibitor |
| *C2* | HM03 | HSPA5 inhibitor | *F7* | ART558 | Polymerase theta inhibitor |
| *C3* | Entinostat | HDAC inhibitor | *F8* | CD532 | AURKA inhibitor |
| *C4* | Ixazomib | Proteasom e inhibitor | *F10* | GSK1520489A | PKMYT1 inhibitor |
| *C5* | Volasertib | PLK1 inhibitor | *G2* | Berzosertib | ATR inhibitor |
| *C6* | Lenvatinib | mTK inhibitor | *G3* | COH34 | PARG inhibitor |
| *C7* | Ethoxysanguina rine | CIP2A inhibitor | *G4* | Azenosertib | WEE1 inhibitor |
| *C8* | Staurosporine | Positive control | *G5* | Thapsigargin | ER stressor |
| *C9* | Lexibulin | Tubulin polymerisat ion inhibitor | *G6* | Eribulin | Microtubule inhibitor |
| *C10* | Ceralasertib | ATR inhibitor | *G7* | Tazemetostat | EZH2 inhibitor |
| *C11* | Olaparib | PARP1/2 inhibitor | *G8* | Camonsertib | ATR inhibitor |
| *C12* | JH-RE-06 | REV1-REV7 inhibitor | *G9* | Carfilzomib | Proteasome inhibitor |
| *D2* | Everolimus | MTOR inhibitor | *G10* | 5Z-7-oxozeaenol | TAK1 inhibitor |
| *D3* | Adavosertib | WEE1 inhibitor | | | |
